# TCR and inflammatory signals tune human MAIT cells to exert specific tissue repair and effector functions

**DOI:** 10.1101/475913

**Authors:** Tianqi Leng, Hossain Delowar Akther, Carl-Philipp Hackstein, Thomas King, Matthias Friedrich, Zoe Christoforidou, Sarah McCuaig, Mastura Neyazi, Carolina V. Arancibia-Cárcamo, Fiona Powrie, Oxford IBD Investigators, Emanuele Marchi, Raphael Sanches Peres, Val Millar, Danie Ebner, Chris Willberg, Paul Klenerman

## Abstract

MAIT cells are an abundant T-cell population enriched in peripheral tissues such as the liver. They are activated both through TCR-dependent and - independent mechanisms. However, the different specific functional responses of MAIT cells to these distinct signals remain elusive. We examined the impact of combinations of TCR-dependent and -independent signals in blood and tissue-derived human MAIT cells. TCR-independent activation of MAIT cells from blood and gut was maximised by extending the panel of cytokines to including TNF-superfamily member TL1A. RNAseq experiments revealed that TCR-dependent and -independent signals drive MAIT cells to exert overlapping and unique effector functions, impacting both host defence and tissue homeostasis. While TCR-triggering alone is insufficient to drive sustained activation, TCR-triggered MAIT cells did show specific enrichment of tissue-repair functions at the level of gene expression, protein production and in *in vitro* assays and these functions were amplified by cytokine costimulation. Taken together, these data indicate the blend of TCR-dependent and -independent signalling to MAIT cells may play a role in controlling the balance between healthy and pathological processes of tissue inflammation and repair.

## INTRODUCTION

Human innate and adaptive immune systems form a critical partnership in immune defence against microorganisms. Recent studies have revealed several types of unconventional T lymphocyte that sit at the bridge between innate and adaptive immunity, including mucosal associated invariant T (MAIT) cells (Godfrey et al., 2015). MAIT cells are abundant in human blood, and enriched most substantially in the liver (Treiner et al., 2003). They are marked by high surface expression of the C-type lectin molecule CD161, and bear the semi-invariant TCR Vα7.2-Jα33/12/20, which restricts them to the evolutionary conserved, non-polymorphic MHC class I-related protein 1, MR1 (Kjer-Nielsen et al., 2012; Reantragoon et al., 2013). MAIT cells recognize microbially derived riboflavin synthesis intermediates presented by MR1 (López-Sagaseta et al., 2013; Ussher et al., 2014). Recently, MR1-tetramers loaded with riboflavin and folate intermediates have been developed, enabling the specific detection and characterization of human and mouse MAIT cells (López-Sagaseta et al., 2013; Rahimpour et al., 2015; Reantragoon et al., 2013).

Despite this specific antigen recognition as an effector T cell, the MR1-TCR signaling alone is insufficient to fully activate MAIT cells (Turtle et al., 2015). In order to achieve sufficient activation, TCR signaling is supported by other co-stimulatory signals such as CD28 and notably by cytokines, for example, interleukin (IL)-18 and IL-12 (Ussher et al., 2013). This is true in mouse cells examined *in vivo* – normal expansion is only seen if ligand is delivered with a TLR-stimulus (Chen et al., 2017).

This behavior has prompted investigation into the responsiveness of MAIT cells to innate signals, including IL-12, IL-18, IL-15, and type I interferons (IFNs) (Sattler et al., 2015; Ussher et al., 2013; Wilgenburg et al., 2016). *In vitro* studies in human cells have shown that cytokines, such as IL-12 and IL-18, can, in combination, activate MAIT cells in a fully TCR-independent manner (Ussher et al., 2013). Cytokine-stimulated CD161^++^CD8^+^ T cells, including MAIT cells, may exert effector functions by secretion of cytokines and upregulation of Granzyme (Gr)B (Billerbeck et al., 2010; Kurioka et al., 2014). We and others recently highlighted a role for MAIT cells in viral infections, where MAIT cell activation was TCR-independent, but dependent on IL-18 in synergy with IL-12, IL-15, and/or type I interferons (IFN-α/β) (Loh et al., 2016; Wilgenburg et al., 2016), with a critical protective role *in vivo* (Wilgenburg et al., 2018). Thus it is clear that MAIT cells can be activated via TCR-dependent and -independent pathways. However, the diversity of functions triggered by different cytokines compared to by TCR signaling has yet to be defined.

The discussion of specific functions of MAIT cells elicited by cytokines is particularly relevant in mucosal tissues, such as the gut, where local signaling may be critical in defining the balance between host defence responses and tolerance. Recent data on IL-17-expressing skin-homing mouse CD8^+^ T cells, an innate-like T cell population which mirrors some critical features of MAIT cells, indicate that they display a tissue repair rather than a pure inflammatory phenotype in response to TCR triggering via commensal associated ligands (formyl peptides restricted by H2-M3) (Linehan et al., 2018). The authors propose that responses to commensals driven by TCR could support a role for such T cells in tissue homeostasis. Indeed this behaviour may extend to more broadly include innate-like T cells restricted by MHC1b molecules, which are evolutionarily ancient (Klenerman and Ogg, 2018).

TNF-like protein 1A (TL1A)/TNF superfamily member 15 (TNFSF15) is a gut-associated proinflammatory cytokine originally characterized in a screen for TNF-α homologous molecules. It is expressed by activated T cells, dendritic cells, and monocytes, and signals through death receptor-3 (DR3) (Meylan et al., 2008; Migone et al., 2002; Shih et al., 2009). TL1A is particularly relevant as it has previously been described to activate a subset of CD4+ memory T cells expressing IL-18Rα and DR3 (Gudjonsson et al., 2015). More specifically, it has been shown to increase production of IFN-γ and TNF-α by CD161^+^CD4^+^ T cells in the presence of anti-CD3 or IL-12+IL-18 (Jin et al., 2012). This may be relevant to MAIT cell functions as we have previously identified a phenotypic, functional, and transcriptional program shared by CD161-expressing cells (Fergusson et al., 2014).

Here, we addressed how TCR-dependent and -independent signals synergize and drive the activation of *in vitro* blood- and gut-derived MAIT cells. We find that IL-12 and IL-18, in synergy with TCR triggering, promotes the activation of MAIT cells in a dose-dependent manner and that TL1A also potently induces specific MAIT cell functions. Interestingly, triggering with TCR alone or supported by cytokines drives a set of functions linked to a tissue repair gene expression signature, accompanied by relevant protein expression and function. Overall, our data provide insight into the precise nature of TCR- and cytokine-mediated human MAIT cell activation, characterized by a broad range of effector functions including not only emergency host defense but also maintaining ongoing homeostasis. This feature may be also relevant to other innate-like T-cell subsets found at barrier sites in humans.

## RESULTS

### TL1A and IL-15 enhance effector functions of human MAIT cells

To explore the full impact of cytokine triggering of MAIT cells we first examined extended combinatorial signaling. The ability of TL1A and IL-15 to promote T cell activation in the presence of a suboptimal IL-12 and IL-18 trigger has been shown in the CD161-expressing CD4^+^ T cells (Cohavy et al., 2011; Jin et al., 2012; Sattler et al., 2015, 2009)(Gudjonsson et al., 2015). We therefore addressed whether MAIT cells possess similar responsiveness.

TL1A triggered MAIT cell activation (as judged by expression of IFN-γ, TNF-α, and Granzyme (Gr) B) in combination with sub-optimal doses of IL12, IL-18 and IL-15 in a dose-dependent manner **(Fig. 1A-C)**. Expression of IFN-γ, TNF-α, and Granzyme (Gr) B by stimulated MAIT cells from a representative donor is shown in **Fig.1D**. IL-15 (50ng/ml), TL1A (100ng/ml), or the combination of both markedly promoted IL-12 (2ng/ml) /IL-18 (50ng/ml) -induced MAIT cell activation, measured by MAIT cell expression of IFN-γ, TNF-α, GrB and CD69 **(Fig. 1E-H)**.

**Figure 1.**
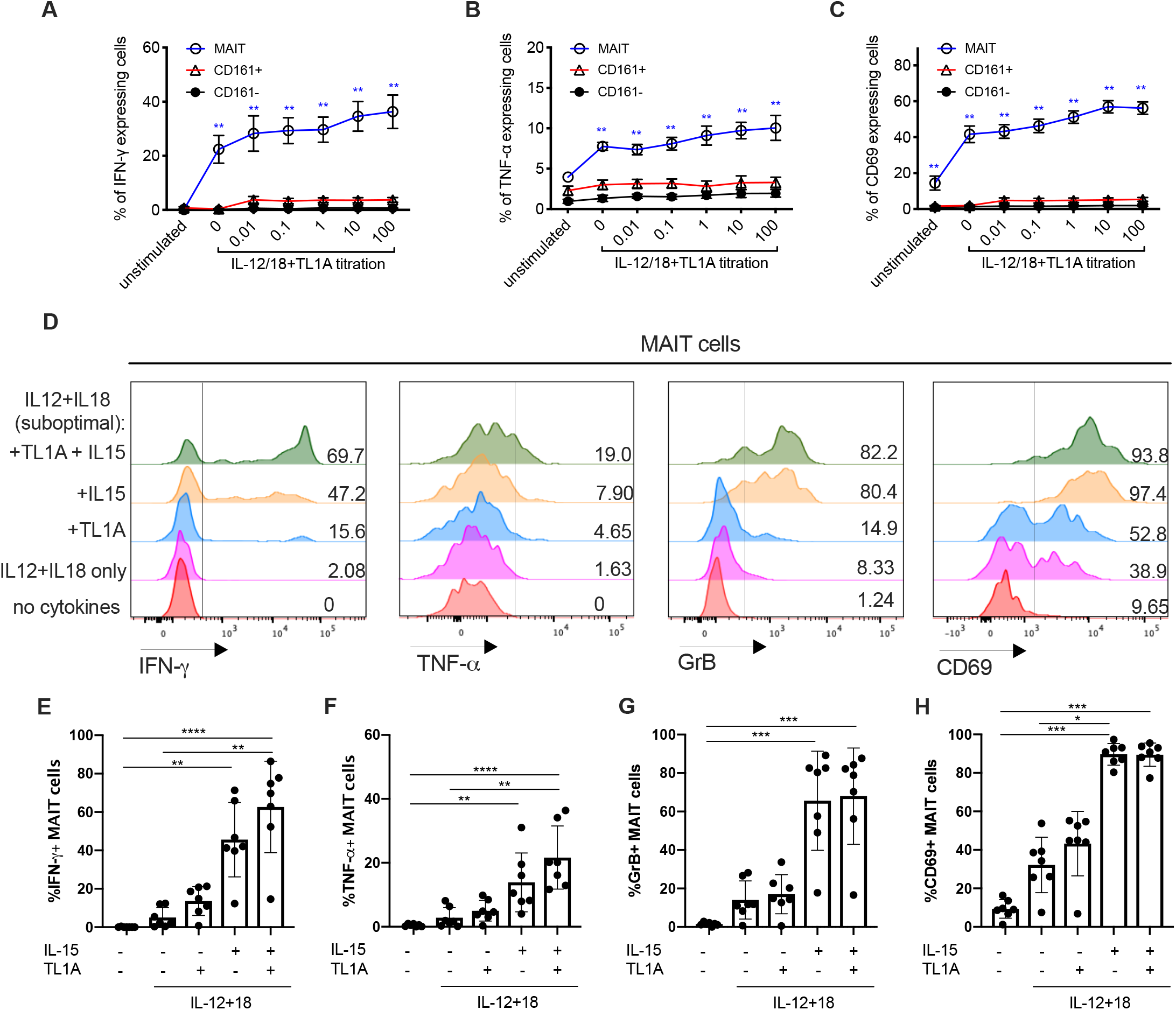
TL1A enhances the activation of MAIT cells suboptimally stimulated with IL-12 and IL-18. CD8+ T cells were enriched from healthy PBMCs and stimulated overnight with different combinations of cytokines: IL-12 used at 2ng/mL, IL-18 at 50ng/mL, IL-15 at 25ng/mL, and TL1A at 0.01-100ng/mL as indicated. **(A-C)** Proportions of CD8+ MAIT/ CD161+ or CD161-cells producing IFN-γ (A), TNF-α (B) or CD69 (C) following overnight stimulation with suboptimal concentrations of IL-12 and IL-18, plus varying concentrations of TL1A. **(D)** Representative histograms showing the expression of IFN-γ, TNF-α, Grb and CD69 by MAIT cells after stimulation with different combinations of cytokines. **(E-H)** Frequency of MAIT cells expressing IFN-γ (E), TNF-α (F), Grb (G) and CD69 (H) upon stimulation with the indicated cytokines. Data were acquired from seven donors in 2-3 experiments. Differences between the conditions were analysed by Friedman tests with Dunn’s multiple comparison tests. *p<0.05, **p<0.01, ***p<0.001, ****p<0.0001.

Overall, we found that TL1A and IL-15, individually increased MAIT cell expression of IFN-γ and TNF-α, and upregulated GrB and CD69 expression. IL-15 was more potent than TL1A when added singly to the IL-12+IL-18 culture, but the peak level of MAIT cell activation was achieved by a combination of both cytokines in the presence of IL-12 and IL-18 – this combination was therefore used in downstream experiments. Of note, TL1A and IL-15 alone do not promote MAIT cell effector functions and have only a limited effect on CD161+ and CD161-CD8+ T-cells **(SFig. 1)**

### MAIT cells respond to combinations of cytokines and TCR triggering and enhance effector functions in a dose-dependent manner

Studies have addressed the hypo-responsiveness of CD8^+^ MAIT cells to anti-CD3 by comparing their ability to proliferate and produce cytokines *in vitro* to their CD161^−^ CD8^+^ counterparts (Turtle et al., 2015). We hypothesized that this could be due to a lack of complementary inflammatory signals. Thus, we first asked how TCR and cytokine signaling combined **(Fig. 2)**. For these experiments we used the optimised MR-1 ligand 5-OP-RU and also compared this to anti-CD3/28 bead stimulations.

**Figure 2.**
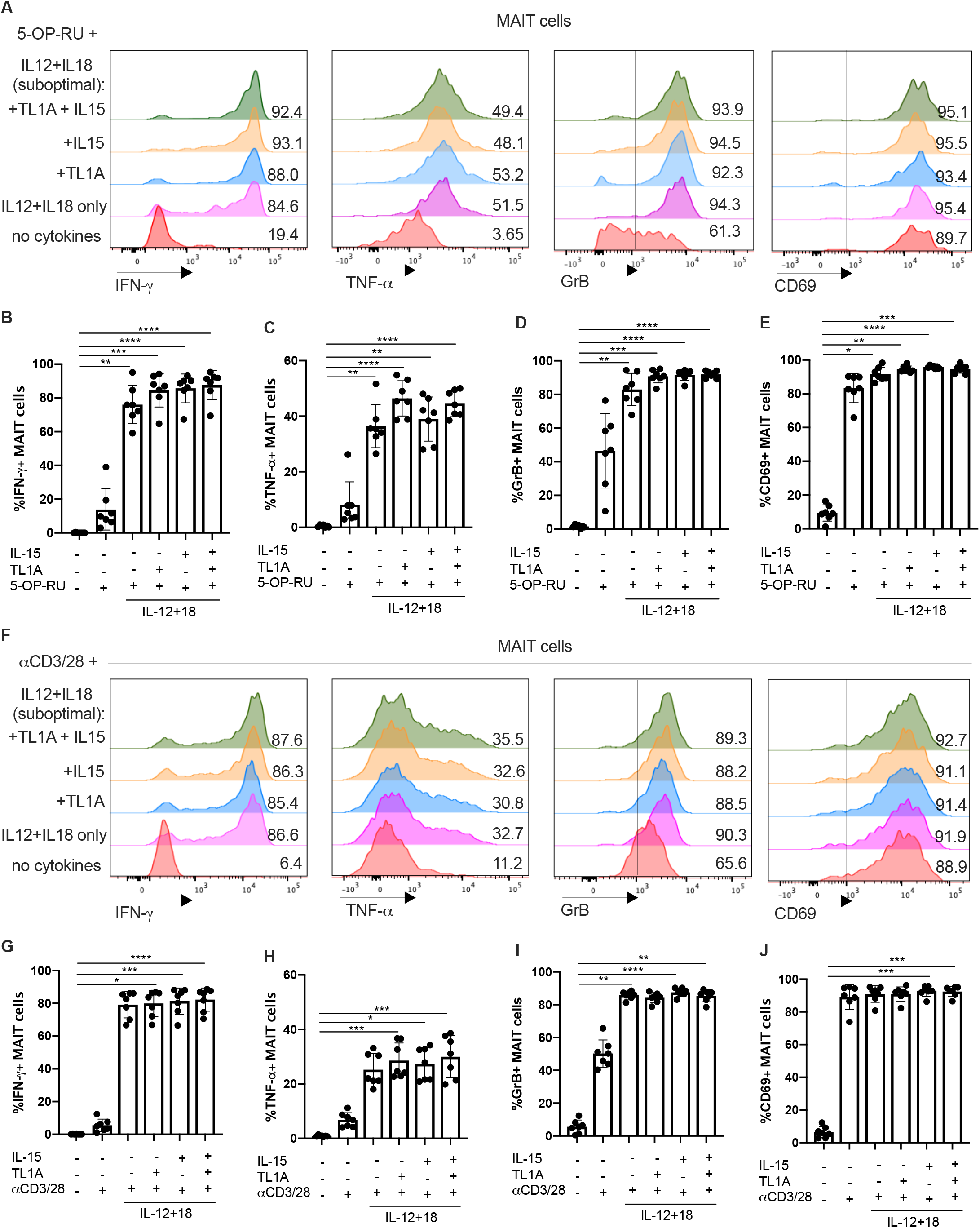
TCR and cytokine signalling combine to promote MAIT cell effector functions. MACS-enriched CD8 T-cells from the blood were cultured overnight in the presence of the indicated cytokines together with the THP1 cell pulsed with DMSO or the MAIT-antigen 5-OP-RU (A-E) or with αCD3/28 beads (F-J). **(A)** Representative histograms showing the expression of IFN-γ, TNF-α, Grb and CD69 by MAIT cells after stimulation with different cytokines in the presence of 5-OP-RU. **(B-E)** Frequency of MAIT cells expressing IFN-γ (B), TNF-α (C), Grb (D) or CD69 (E) upon stimulation with the indicated cytokines. **(F)** Representative histograms showing the expression of IFN-γ, TNF -α, Grb and CD69 by MAIT cells after stimulation with different cytokines in the presence of 5-OP-RU. **(G-J)** Frequency of MAIT cells expressing IFN-γ (G), TNF-α (H), Grb (I) or CD69 (J) upon stimulation with the indicated cytokines. Data were acquired from seven donors in two experiments. Differences between the conditions were analysed by Friedman tests with Dunn’s multiple comparison tests. *p<0.05, **p<0.01, ***p<0.001, ****p<0.001

Enriched CD8+ T cells were stimulated by suboptimal concentrations of IL-12 (2ng/ml) and IL-18 (50ng/ml), +/− TL1A and/or IL-15 in combination with 5 OP-RU **(Fig. 2A-E)**. TCR signaling via 5-OP-RU had a profound synergy with added cytokines – with the major impact from IL-12 and IL-18 – as measured by release of IFNg and TNFa, upregulation of GrB and CD69 (representative histograms **Fig. 2A**, combined data **Figs. 2B-E**).

We repeated these protocols using anti-CD3/CD28 beads as the TCR trigger (**Fig. 2F-J**). The frequency of MAIT cells that responded to anti-CD3/CD28 by producing IFN-γ or TNF-α positively correlated to the bead-to-cell ratio **(SFig. 2A and B)**. Very similar data were obtained to those using the 5-OP-RU trigger above (histograms **Fig. 2F**, combined data **Figs. 2G-J**). Again in these combined TCR-cytokine stimulation studies IFN-γ expression correlated with that of CD161, consistent with previous findings (Fergusson et al., 2015, 2014), and TL1A and IL-15 had no impact individually (**SFig. 2C**).

### Gut-derived MAIT cells also respond to a range of cytokine and TCR stimulations and show evidence of activation *in vivo*

We next examined MAIT cells in the gut to address the role of signal integration in tissue-derived MAIT cells. A gating strategy for MAIT cells on a representative sample is shown (**SFig. 3A, C**). An additional stain for γ*δ* T cells was included to show that the number of γ*δ* TCR-bearing CD8^+^ T cells is minimal. For all subsequent experiments, the γ*δ* T cell population was excluded from analysis using a dump channel. Colonic lymphocytes were isolated from uninvolved mucosa from colorectal cancer resection tissue.

MAIT cell activation was assessed by their expression of IFN-γ, TNF-α and GrB (**Fig. 3A-F**), in response to TCR and cytokine triggers alone or in combination as above. TCR and sub-optimal IL-12/IL-18 triggers synergized strongly, and maximal activation was seen using combined stimulations including IL-15 and TL1A. This pattern was clearly seen in blood (**Fig. 3 A-C**) and in gut-derived cells (**Fig 3D-F**), although the absolute magnitude of the stimulation was lower in the latter. We confirmed that this lower responsiveness was not due to discrete preparation conditions (**SFig. 3B**).

**Figure 3.**
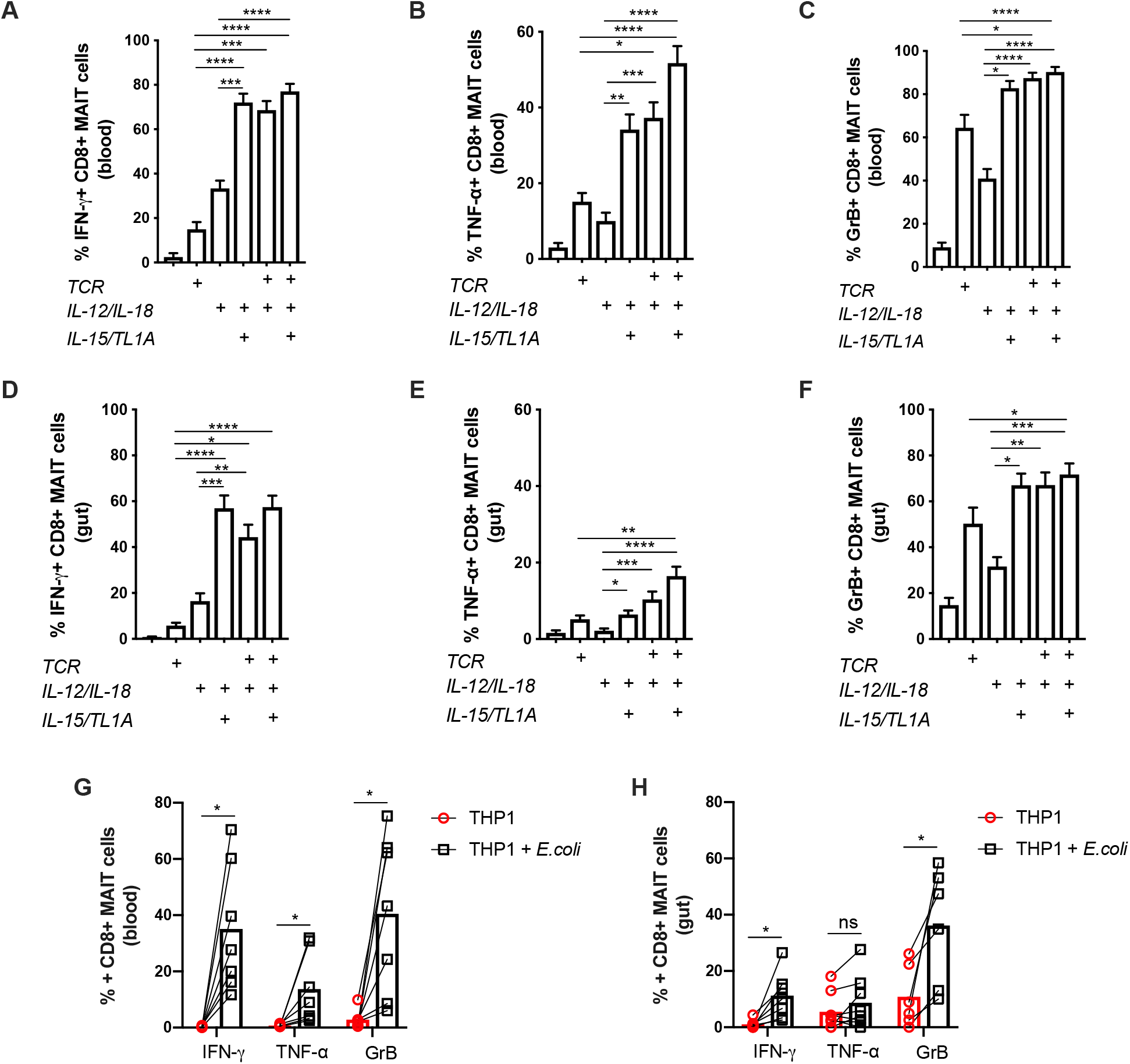
Gut-derived MAIT cells show a similar but dampened response pattern towards innate and adaptive stimuli compared to their blood-derived counterparts. Representative plots showing the percentage of cell positive for the indicated effector molecules as a proportion of CD8+ MAIT cells. **(A-C)** Proportions of blood-derived (n=32) CD8+ MAIT cells producing IFN-γ (A), TNF-α (B) or GrB (C) following overnight stimulation with combinations of subpotimal concentrations of IL-12 and IL-18, TL1A and αCD3/28 beads as indicated. **(D-F)** Proportions of gut-derived (n=13) CD8+ MAIT cells producing IFN-γ (D), TNF-α (E) or GrB (F) stimulated the same way as in (A-C). **(G, H)** Expression of IFN-γ, TNF-α and GrB by blood-(G, n=&) or gut-derived (H, n=6) CD8+ MAIT cells 20h after co-culture with THP1 cells alone or THP1s incubated with 25 fixed *E.coli* bacteria per cell. Data were acquired from multiple donors as indicated in 3-5 experiments. Differences between the conditions were analysed by Friedman tests with Dunn’s multiple comparison tests (A-F), 2-way ANOVA (G) or Wilcoxon tests (H). *p<0.05, **p<0.01, ***p<0.001, ****p<0.001

Since these data were observed using TCR beads, we repeated these experiments with a more physiological ligand derived from *E coli*, using an established, MR1-dependent protocol (**Fig. 3G, H, SFig. 3D**). Again gut-derived cells showed a similar overall pattern of responsiveness to blood-derived cells, with a blunted magnitude.

We next sought evidence that human MAIT cells at a barrier surface exposed to commensal cues would show some features of ongoing activation. MAIT cells in blood are low in GrB but rapidly upregulate this in response to TCR and cytokine signals. When we examined MAIT cells from the gut we observed a consistent upregulation of GrB (**Fig. 4A**). Chronic TCR stimulation e.g. in viral infections such as LCMV, HCV and HIV can lead to upregulation of both CD39 and PD-1 on CD8+ T cells (Gupta et al., 2015). PD1 and CD39 were both significantly elevated on MAIT cells derived from the control gut (**Fig. 4B, C**), and consistent findings were made by measuring MAIT cell PD-1 and CD39 geometric mean of fluorescence intensity (**Fig. 4D, E**). We then examined the co-expression of these two markers and found a significantly elevated proportion of CD39^+^ PD-1^+^ MAIT cells in the control mucosa than the blood (**Fig. 4F, G**). Taken together, these data suggest ongoing, sustained triggering of MAIT cells in a healthy gut, with phenotypic and functional adaptation. They also indicate that gut-derived MAIT cells respond to both TCR and cytokine signals in an overall pattern similar to that seen with blood-derived cells.

**Figure 4.**
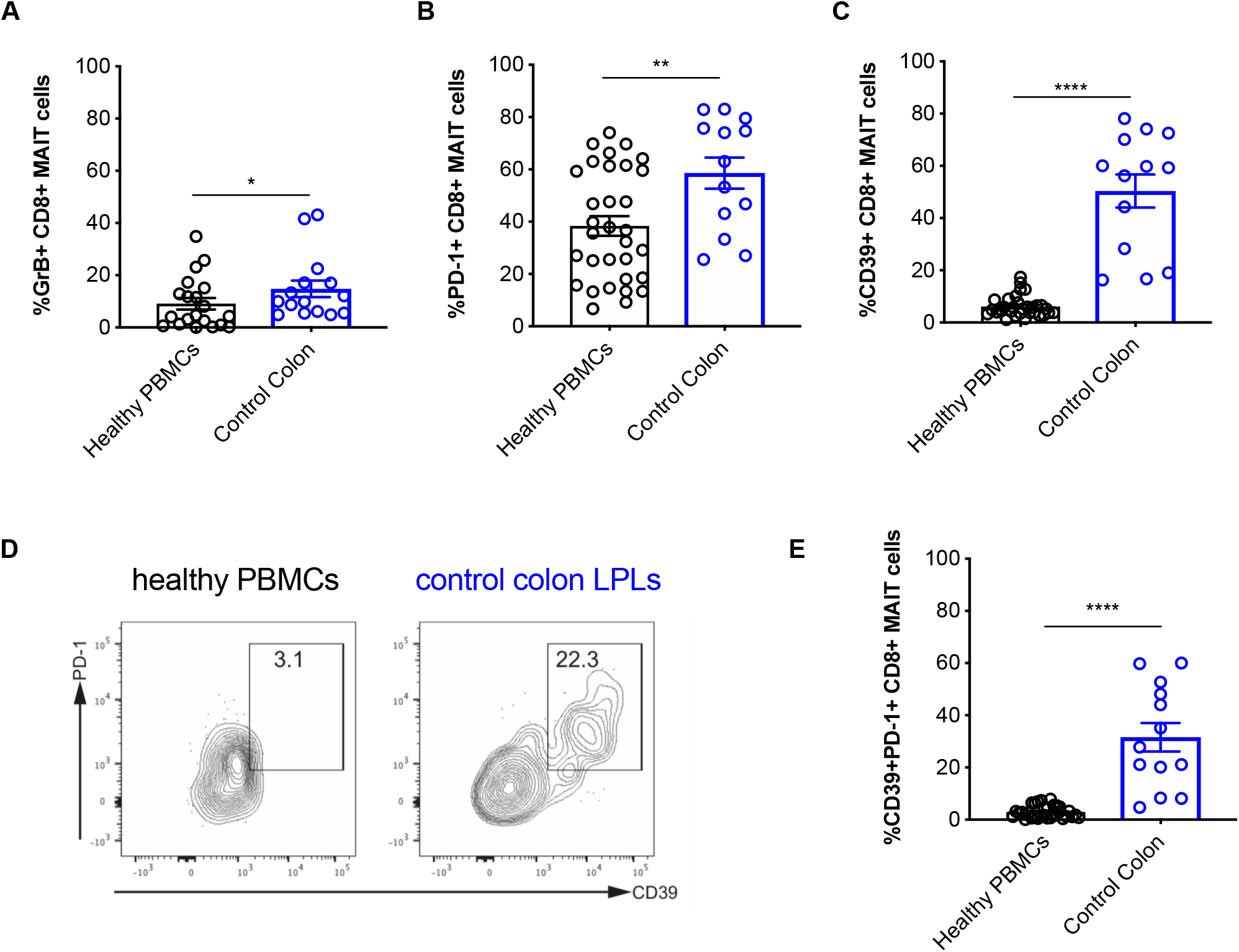
MAIT cells derived from the healthy gut display increased expression of PD-1 and CD39. Healthy PBMC’s or LPLs extracted from healthy gut tissues were analysed by flow cytometry. **(A-C)** Proportion of MAIT cells from the blood and tissue expressing GrB, PD-1 or CD39. **(D)** Representative plots showing the co-expression of PD-1 and CD39 in MAIT cells derived from the blood and the gut. **(E)** Proportion of PD-1+CD39+ MAIT cells from the blood and gut. Data were acquired from 13 (LPLs) or 33 (PBMCs) donors in 13 experiments. Differences between conditions were analysed by Mann-Whitney tests. *p<0.05, **p<0.01, ****p<0.001

### MAIT cells possess distinct transcriptional signatures upon activation by TCR or cytokines

To further explore the full breadth of effector functions of MAIT cells elicited by the TCR or cytokine signals, we utilized RNAseq to characterize transcriptional profiles of MAIT cells under different treatments, including the TCR (anti-CD3/CD28, labeled here as “T”), Cytokines (IL-12/IL-15/IL-18/TL1A, labeled here as “C”), and a combination thereof (labeled here as “TC”). Transcriptional profiles of differentially stimulated MAIT cells were compared to those of untreated (UT) cells. TCR beads were used at 1:1 bead-to-cell ratio, and cytokines were used at the concentrations optimised above. To confirm activation, MAIT cells from the same donors were also examined for their release of IFN-γ, TNF-α and expression of GrB in response to the same stimulations (**SFig. 4A-C**).

The mRNA levels of 132, 1124, and 1375 genes were significantly modulated (p < 0.01, |fold change| > 4, false discovery rate (FDR) < 0.05, including upregulation and downregulation) by TCR, cytokines, or combined TCR and cytokine stimulation, respectively. Venn diagrams highlight the overlapping and unique transcriptional signatures elicited by these 3 different stimulations (**Fig. 5A-C, Supplementary Table 1, Sheet_1)**. We found that stimulating MAIT cells with TCR beads and/or cytokines resulted in the significant alteration of 89 common mRNA transcripts in MAIT cells, including 88 upregulated genes (**Fig. 5B, Supplementary Table 1, Sheet_2**), and 1 downregulated gene (**Fig. 5C**). Gene Ontology (GO) enrichment analysis on these common 88 upregulated genes by MAIT cells predicted that they are involved in cytokine production and signalling including, of relevance, IL-12-mediated signaling (IL23R, EBI3, IL2RA, RELB, NFKB1, NFKB2, CCL3), and TNF signaling (TNF, NFKB1, NFKBIA).

**Figure 5.**
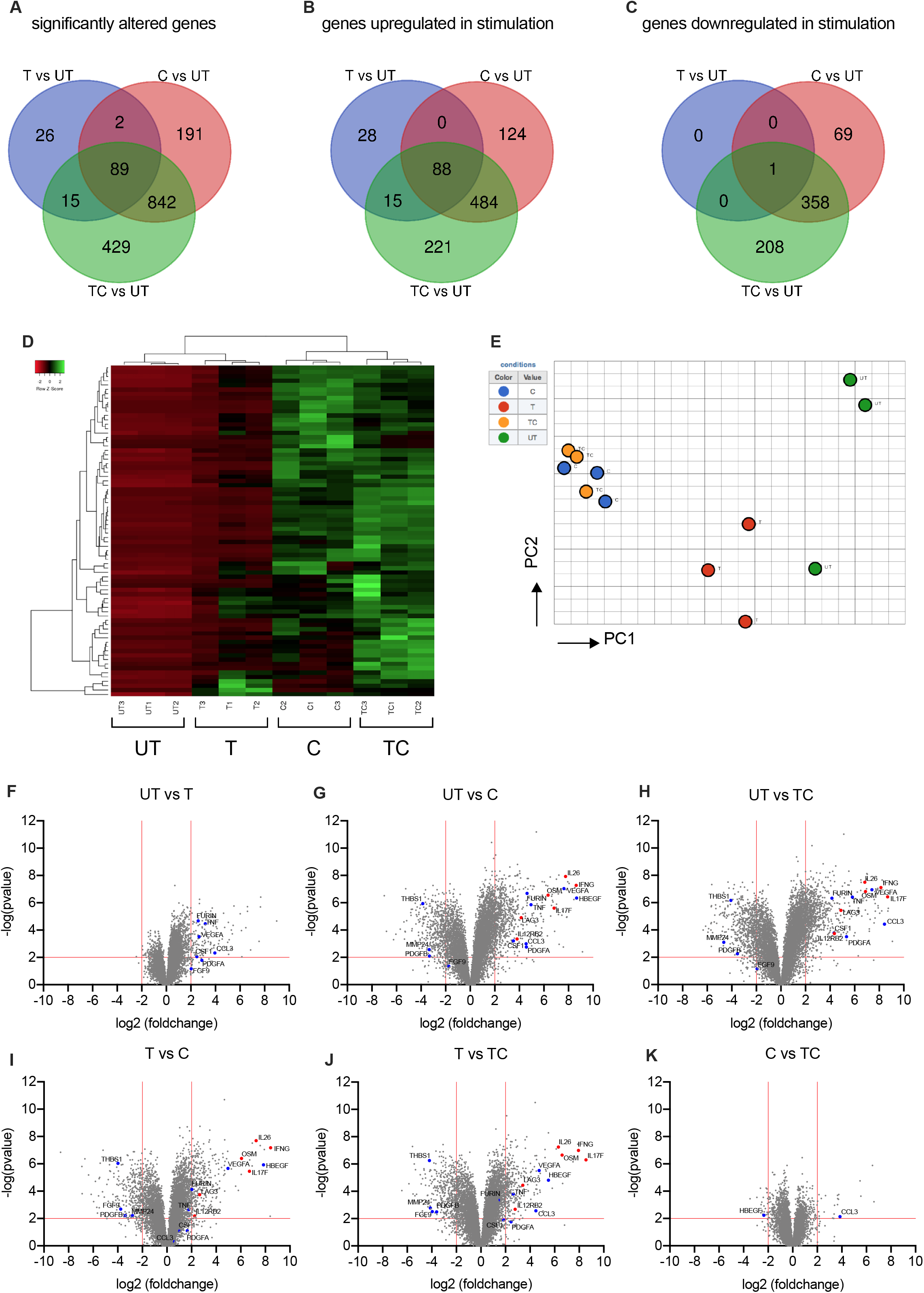
TCR-and Cytokine-activated MAIT cells possess distinct transcriptional profiles. **(A-C)** Venn diagrams showing genes that are significantly differentially modulated (p<0.05, fold change>4) in TCR (T)-/Cytokines(C)-/TCR+Cytokines (TC)-treated CD8+MAIT cells compared with untreated (UT) MAIT cells of three healthy individuals. The cytokine stimulation consisted of a cocktail of 4 cytokines: IL-12 (2 ng/ml), IL-18 (50 ng/ml), IL-15 (25 ng/ml), and TL1A (100 ng/ml). Genes with significantly altered expression levels are further divided into two sets, those are that are upregulated upon stimulation (B), and downregulated (C) upon stimulation. **(D)** Heatmap showing 1,594 significantly (p < 0.05, fold change >4) differentially expressed transcripts between T/C/TC-stimulated and UT CD8+MAIT cells amongst the same three healthy individuals. **(E)** Visualization of the CD8+MAIT cell transcripts elicited by differential stimulations in the subspace of the first principle components (PC). Each coloured circle represents a sample and they are colour-coded in accordance with the conditions with which cells were stimulated, as illustrated on the right-hand side of the graph. **(F-K)** Volcano plots to visualize differentially expressed transcriptional profiles of activated CD8+MAIT cells stimulated with in different ways. Each point represents a single gene, and genes expressed at significantly higher or lower between the compared conditions are depicted on the upper-right or respectively in the upper-left corner of each plot. Genes discussed in the text are highlighted in blue (tissue-repair associated) or in red (inflammation-associated). Data were acquired from three donors in one experiment.

Analysis of the other genes unique to TCR and cytokines indicate diverse physiological functions that these two signals induce in MAIT cells (**Fig. 5A**). Among 1594 modulated genes (**Supplementary Table 2**), 960 (60.2%) were upregulated and the rest were downregulated (**Fig. 5A, B**). MAIT cells stimulated via TCR and cytokines shared 572 upregulated genes with their counterparts only stimulated with cytokines. They constitute 82.1% of upregulated gene transcripts elicited by cytokines, and 70.8% elicited by TCR beads and cytokines (**Fig. 5B**).

The 1594 genes with significantly altered expression levels between conditions were then plotted in a heat map according to their normalized expressions using average linkage hierarchical clustering (**Fig. 5D**). We also performed a principal component analysis (PCA) using the first two principal components of the 1594 mRNA transcripts and visualized the correlation of the transcriptional profiles of differentially stimulated MAIT cells (**Fig. 5E**). Clustering of C- and TC-conditions confirmed that cytokine stimulation at this time-point had a dominant impact on MAIT cell activation. However, the finding of 221 upregulated genes (**Fig. 5B**) and 208 downregulated genes (**Fig. 5C**) unique to TC stimulation also suggests a strong synergy between the TCR signaling and cytokines to further drive MAIT cell activation and promote their effector functions. Furthermore, the PCA analysis indicates that a TCR stimulus alone can trigger a pronounced level of activation, as the T clustering was clearly separate from UT.

We next analyzed volcano plots that show differentially expressed transcriptional profiles of stimulated MAIT cells compared to their unstimulated counterparts (**Fig. 5F-H**) and comparing between different stimulations (**Fig. 5I-K**). Overall a more limited number of genes were significantly altered by a single dose of TCR stimulation, after filtering for transcripts with p value <0.01 and fold change > 4 (**Fig. 5F**), compared to more dynamic transcriptional profiles seen following stimulation with cytokines (**Fig. 5G, H**). The additional transcriptional impact of cytokines on top of TCR stimulation (T vs TC) is shown in **Fig. 5J**.

To confirm these findings from RNAseq, we firstly used qPCR to validate 3 of the most highly upregulated genes - IL-26, oncostatin M (OSM), and Heparin binding early growth factor (HBEGF) **(Fig 5.G, H)** - on RNA samples extracted from activated MAIT cells (**SFig. 4E-G)**. In all 3 cases we demonstrate a similar pattern of responsiveness by an independent method. Overall these data indicated that a very wide range of responses can be generated by MAIT cells in response to TCR and cytokine triggers and that the pattern of these differ between the triggers used – we therefore went on to explore the significance of this in more depth.

### Transcriptional signatures of activated MAIT cells predict not only antimicrobial but also tissue repair functions

Given the broad range of responses seen after TCR and cytokine mediated activation we speculated that functions of MAIT cells extended beyond conventional antimicrobial functions.The recent discovery of a skin-homing Tc17 subset in mouse responsive to commensal ligands has shed light on a unique form of adaptive immunity where antimicrobial functions and tissue repair are coupled within the same subset of unconventional T cells (Harrison et al., 2019; Linehan et al., 2018). These commensal-specific T cells elicited a tissue repair signature and accelerated wound closure, in addition to promoting protection to pathogens. MAIT cells are commensal responsive and similarly have been associated with a type-17 phenotype (Billerbeck et al., 2010; Dusseaux et al., 2011). Therefore, we investigated functional overlap using a genomic comparison between activated human MAIT cells and mouse skin-homing Tc17 cells.

Firstly we examined the volcano plots in Fig 5.F-K and annotated the genes from the tissue-repair gene-list used in the study of Linehan and colleagues **(Supplementary Table 3)**. The genes on these plots **(Fig 5.F-K)** are colour-coded according whether they associated with a pro-inflammatory and anti-microbial response, as has been classically associated with MAIT cells (red) or a tissue-repair signature (blue). Substantial numbers of genes linked with the tissue-repair signature were observed, including genes such as Furin, TNF, CSF1, CCL3 and various growth factors.

Next genes significantly differentially expressed compared to unstimulated MAIT cells were identified from T, C, and TC –stimulated MAIT cells and statistically compared in aggregate to the tissue repair gene data set (Linehan et al., 2018). Remarkably, gene set enrichment analysis demonstrated significant enrichment (p<0.0002) of these tissue repair related genes in MAIT cells stimulated by TCR ± cytokines (**Fig. 6A, B**), but not by cytokines alone (**Fig. 6C**). The significant leading edge genes from these analyses are indicated in **SFig. 5A-B**. These data suggest that TCR triggering by MAIT cells may be important in driving a tissue repair program.

**Figure 6.**
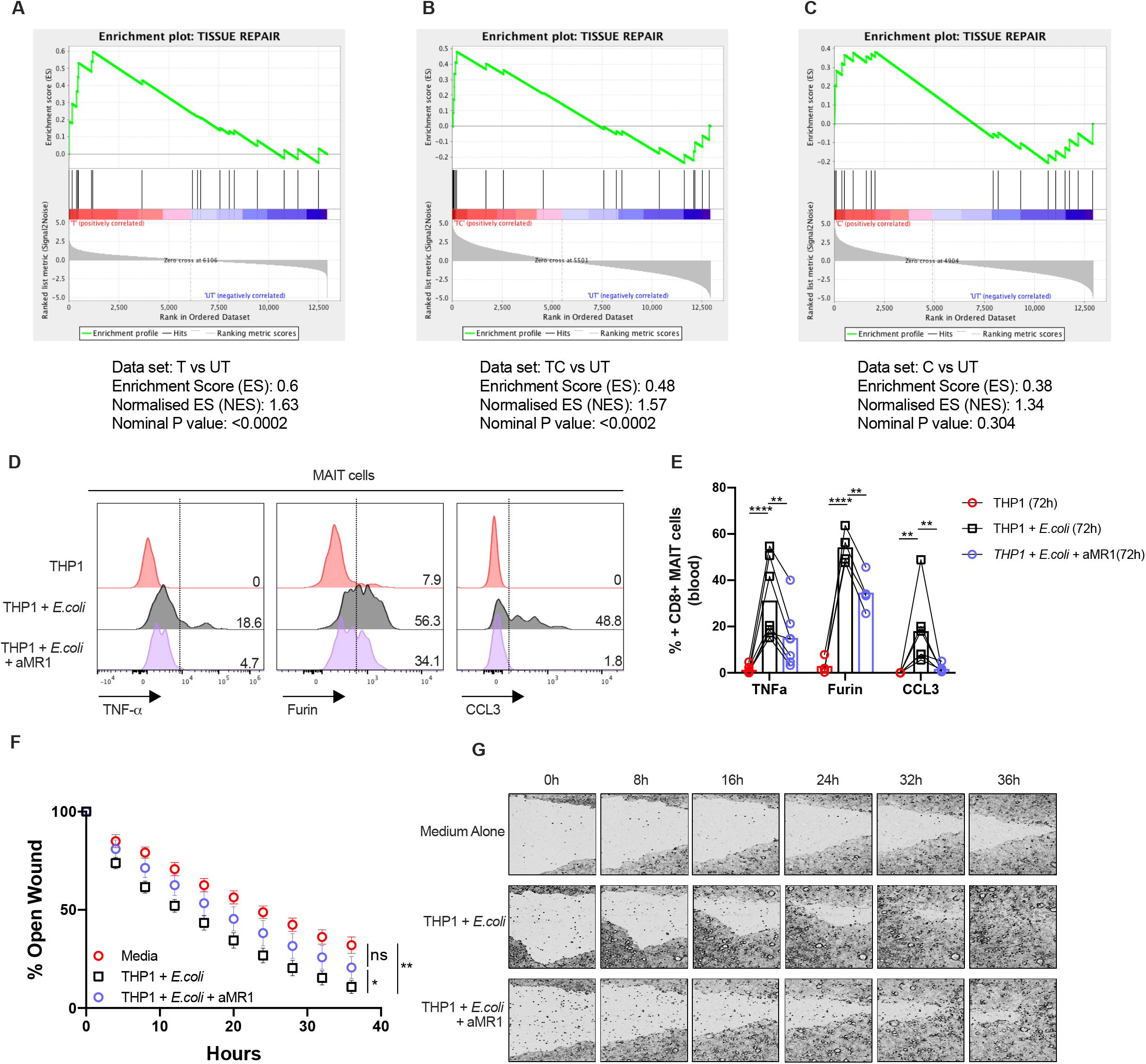
TCR mediated activation of MAIT cells leads to the expression of tissue repair associated molecules and accelerates wound healing. **(A-C)** Gene set enrichment summary plots for cytokine-stimulated (C), TCR-stimulated (T), or TCR+cytokine (TC)-stimulated sorted MAIT cell versus unstimulated (UT) cell-ranked genes. Non-significant for C vs UT, NES = 1.63, p < 0.0002 for T vs UT, NES = 1.57, p < 0.0002 for TC vs UT. Data were acquired from three donors in one experiment. **(D)** Flow cytometry analysis of the expression of TNF-α, Furin and CCL3 by CD161++/MAIT CD8+ T cells in response to fixed *E.coli* presented by THP1 cells in the presence or absence of an anti-MR1 (aMR1) blocking antibody at 72h timepoint. **(E)** Statistical analysis of the expression of the effector molecules shown in (D). **(F)** Caco2 cells were grown to confluency and scratched with a Wound-Maker to perform *in vitro* wound healing assays. Cells were supplemented with different supernatants collected from 72h co-cultures of enriched CD8 T-cells with *E.coli* loaded THP1s in the presence or absence of αMR1, as indicated. The open wound areas were quantified as percentage of the initial wound size in the Caco2 cultures. Data points are Mean +/− SEM and were acquired from five biological replicates in two experiments. **(G)** Representative pictures of the closure of the wounds in Caco2 cultures treated as in (F) were assessed with time-lapse imaging over a time-course of 36h. Data were acquired from seven donors in three experiments. Differences between the conditions were analysed by two-way ANOVA. ns = not significant, *p<0.05, **p<0.01, ****p<0.001.

To test these findings further we analysed the release of 3 of these genes from the tissue-repair signature using flow cytometry. Triggering of MAIT cells by *E coli* led to the production of TNF, Furin and CCL3 in a manner which was fully or partially blockable by anti-MR1 at 20hrs and 72 hrs respectively **(Fig. 6D-E; SFig. C-D)**. We also validated upregulation of GM-CSF (CSF2), a tissue-repair associated gene not upregulated at earlier time points, again most evident in extended cultures and triggered in an MR1-dependent fashion **(SFig. 5E-F)**.

To assess this further functionally we tested the impact of supernatants derived following stimulation of MAIT-cells in wound healing assays in vitro, taking advantage of the scratch assay in Caco2 cells (see methods (Povoleri et al., 2018)). We observed wound closure in this system which was accelerated by MAIT cell supernatants and partially blocked by anti-MR1 **(Fig. 6F-G)**. This was most evident at later timepoints (e.g. 24-36hrs; **Fig 6.G**).

## DISCUSSION

CD161-expressing human T lymphocytes possess shared transcriptional and functional phenotypes, and their enhanced innate ability to respond to inflammatory cues has been investigated with transcriptomic approaches (Fergusson et al., 2014). MAIT cells comprise a large proportion of these CD161-expressing T cells, and have previously been described to show limited responses to conventional TCR signals although combinatorial signalling can markedly augment this in vitro and also in vivo (Slichter et al., 2016; Turtle et al., 2015)(see also accompanying paper by Hinks and colleagues (Hinks et al., 2018)). On the other hand, MAIT cells can respond in a fully TCR-independent manner, via cytokine signalling, and such behaviour can also trigger protection in vivo (Wilgenburg et al., 2018). The functional consequences of TCR-dependent vs TCR-independent activation of MAIT cells have not been fully defined - thus to further dissect the differential signals that promote and sustain MAIT cell effector function, is of central importance in defining the role of MAIT cells in both health and disease. Here we probed the contribution of innate and adaptive signals to MAIT cell activation in the blood and gut, describing and segregating novel functions of MAIT cells following activation.

We defined MAIT cells in both human blood and gut as CD161^++^Vα7.2^+^CD8^+^CD4^−^ CD3^+^ T cells, a common approach used by a number of studies to dissect MAIT cell effector function elicited by cytokines or the TCR(Kurioka et al., 2014, 2017; Sattler et al., 2015; Ussher et al., 2013; Wilgenburg et al., 2016). MR1 tetramers, combined with CD161 staining, identify MR1-restricted MAIT cells unequivocally, but these reagents only became available at the end of this study. We also stained freshly isolated colonic lymphocytes using the MR1 tetramer, with Va7.2 and CD161 antibodies on the same panel. These provided similar estimates of frequency (as they commonly do in blood) – an example is shown (**SFig. 3C**).

Responses following a TCR stimulation of MAIT cells and conventional CD8 T cells differ in magnitude and also in quality. We and others (Slichter et al., 2016) have initially explored this using bead based protocols – this allowed a direct comparison with non-MAIT populations and simplified some downstream sorting procedures. Although CD3/28 beads do not represent a true physiologic TCR stimulus, here we have confirmed the bead-based data using 5-OP-RU, an optimised MR1 ligand, and also E coli stimulation, which represents a more physiologic stimulus. In each case there is clearly activation in response to TCR-triggering, but this is markedly amplified and sustained through combinatorial signals via cytokines. These broad features were recapitulated in MAIT cells derived from the human gut.

Resting MAIT cells express a higher level of IL-18Rα on the cell surface, compared to their CD161^++^Vα7.2^−^ or CD161- counterparts (Fergusson et al., 2014; Ussher et al., 2013). However, we found that IL-18 had a very limited role in enhancing MAIT cell activation in the presence of TCR triggering at 20 hours. Studies also showed that other human lymphocytes, including NK cells and T cells, did not produce IFN-γ when singly treated with IL-18 (Jampel et al., 2009; Tominaga et al., 2000). The IL-18 receptor is a heterodimer comprising IL-18Rα (IL-1Rrp) and IL-18Rβ (AcPL) chains. Both subunits are markedly upregulated in cytokine-activated NK and T lymphocytes, including IL-12 and IFN-α (Sareneva et al., 2000), but not by TCR signals. Thus, the observed low response of MAIT cells to TCR+IL-18 stimulation maybe due to insufficient expression of IL-18Rβ, which is important for downstream signalling (Cheung et al., 2005). A signal combining IL-12 and IL-18 has been described to activate a range of lymphocytes, including Th1 cells, B cells, NK cells, and more recently, MAIT cells, independently from the TCR. Titrating TCR signals into a suboptimal IL-12+IL-18 coculture restored effector function of MAIT cells (SFig 2). IL-12 and TCR synergy likely results from IL-12-mediated phosphorylation of Lck, a key player in the TCR signaling (Vacaflores et al., 2016). To summarize, our work provides evidence that inflammatory cytokines and the TCR signal synergise to enhance MAIT cell effector function.

Our data also show evidence of TL1A-mediated MAIT cell activation. In addition to its pro-inflammatory and costimulatory role in the human blood, TL1A is a gut-associated cytokine, and has been linked to IBD (Jin et al., 2012; Shih et al., 2009). Patients with IBD have higher levels of DR3 and TL1A expression in their mucosal T cells and macrophages (Bamias et al., 2003; Prehn et al., 2004). We highlight that TL1A enhanced the effector function of gut MAIT cells in the presence of a suboptimal dose of IL-12+IL-18, suggesting that TL1A may contribute to the amplification of inflammatory responses. Interestingly we found that TL1A impacted on MAIT cell TNF-α production in response to inflammatory stimuli but blockade did not impair responsiveness to bacteria. Thus, TL1A blockade *in vivo* could potentially achieve an anti-inflammatory response while maintaining barrier function. These data confirm and extend the importance of TNF superfamily members in MAIT cell activation – as recently revealed for TNF-α itself in responses to opsonised bacteria at limiting doses (Banki et al., 2019), and suggest a context-specific and APC-dependent role for these signals.

Genes controlling the TCR signaling pathway have shown to be differentially regulated in MAIT cells compared to conventional CD8^+^ T cells (CD161^−^ cells) (Turtle et al., 2015), yet full implications of a partial response by TCR-triggered MAIT cells have been so far unclear. The conserved MAIT TCR consisting of the invariant TCRα chain and a restricted repertoire of β chains allows MAIT cells to specifically recognize bacterial metabolites, which can be derived from either pathogenic or commensal bacteria. Given the abundance of both MAIT cells and commensal bacteria in the human gut (Serriari et al., 2014), we speculated that MAIT cells could tune TCR signals to optimise activation for local responsiveness when encountering commensal-derived antigens and to amplify these signals towards a full inflammatory response in the setting of invasive pathogens. We therefore explored this using RNAseq of MAIT cells stimulated in vitro via TCR-dependent and -independent pathways.

Our RNAseq data reveal both shared and independent response patterns between TCR-dependent and -independent pathways. Notably although the magnitude of change seen in our experiments was greater in the cytokine-stimulated cells, the transcripts associated with TCR triggering alone provided a clear insight into potential function. We linked, using GSEA, the MAIT TCR-driven transcriptional profile with a tissue repair signature from a recent report on IL17+ innate like CD8+ T cells in a murine skin model (Linehan et al., 2018) (later confirmed in the gut (Harrison et al., 2019)). The tissue repair profile in the unconventional (H-2 M3-restricted) mouse skin CD8+ T cells was shown to be linked to encounter with commensal microbes and to impact on cutaneous wound healing. In our model we have shown a functional impact of TCR-dependent microbe-triggered MAIT cells *in vitro* using a monolayer scratch assay.

We validated some of the most critical genes underpinning this signature, such as furin, using separate tools and confirmed that MAIT cells respond by secretion of these critical proteins. Furin plays an important role in tissue repair through its broad proprotein convertase activity leading to activation of proteins like TGFβ and matrix metalloproteinases. T-cell derived furin has been shown to be critical in tissue protection in a transfer colitis model and has impacts on Treg development (Pesu et al., 2008). CCL3 (MIP1a), which is more broadly expressed and has been better studied in T cells has a critical role in macrophage recruitment (DiPietro et al., 1998). Both of these mediators and also TNFα – which has a described role in tissue repair in concert with CCL3 (Li et al., 2016)– were produced by activated MAIT cells in a sustained and TCR-dependent manner. The repair signature was most evident in MAIT cells triggered via their TCR, either with or without cytokines, but there was no statistical enrichment in the cytokine-only stimulation. This is most evident in Fig 5F-K, where some of the key genes are marked. For example, the cytokine-alone stimulus induces many relevant genes but there is a slight abundance of inflammatory/host defence genes upregulated in the C vs T comparison (Fig 5I vs 5F), together with some downregulation of tissue repair genes from the GSEA list (Supplementary Table 3).

Beyond the list of tissue-repair factors identified in the work of Linehan (Linehan et al., 2018) and colleagues and used in the GSEA comparison, other MAIT-cell derived factors were found through the RNAseq study (and validated in independent assays) with potential roles in inflammation and also in tissue homeostasis. For example, IL-26 is part of the IL-20 family of cytokines (including Il-22, also made by MAIT cells) which all strongly impact on epithelial cell function including wound repair (Rutz et al., 2014). Similarly OSM, which is upregulated in the gut during inflammation in vivo (West et al., 2017), has been shown to induce migration of keratinocytes *in vitro* and skin repair *in vivo* (Boniface et al., 2007; Hoffmann et al., 2011), while HBEGF, which was highly expressed by activated MAIT cells has a long track record in tissue regeneration (reviewed in (Dao et al., 2018)). Taken together these data substantially broaden the known functions of MAIT cells and include a range of functions on a spectrum between host defence, inflammation and barrier repair. A very similar set of RNASeq data, functional data and conclusions has been obtained using parallel experiments in human MAIT cells *ex vivo* with a 5-OP-RU trigger and most importantly in an in vivo challenge incorporating a TCR trigger with and without cytokines (Hinks et al., 2018). Thus, overall a reasonable body of data has emerged in parallel which suggests MAIT cells possess tissue repair activity of relevance in barrier defence.

In contrast to their roles in host defence and in inflammation there is yet to be a consensus established around the function of CD8+ T cells in wound healing and tissue repair. The precise functions of different T cell types and subsets are largely unknown and differ from tissue to tissue and between species. While γδ T cells are commonly believed to promote tissue healing processes (Havran and Jameson, 2010), studies around CD4+ Tregs and CD8+ cytotoxic cells have suggested their diverse and distinct roles in wound healing. In the context of bone, fracture healing is accelerated in mice depleted of T and B cells (Toben et al., 2011), whereas other studies have shown that CD4+ Tregs are critical to repair and regenerate several tissues including skin and lung (Julier et al., 2017), and that CD8+ T cells aid skeletal muscle regeneration (Zhang et al., 2014) and wound closure in mice (Linehan et al., 2018), with an important role for IL-18 in such activity (Harrison et al., 2019).

Taking together the data here and those of Hinks (Hinks et al., 2018) we propose a model whereby in the human gut and potentially also in the liver, MAIT cells are continuously exposed to MR1-bound ligand derived from the commensal bacteria present in the microbiome. In the absence of inflammatory signals, this serves only to drive the circumscribed transcriptional signature associated with local homeostatic function. Consistent with this, gut MAIT cells show evidence of persistent TCR-driven signalling (upregulation of CD39 and PD1) together with downregulation of function compared to blood-derived cells. During loss of bowel integrity associated for example with inflammatory bowel disease, MAIT cells have been shown to be further activated in some studies (Serriari et al., 2014), but whether this is a response to tissue injury or they play a causative role is yet to be defined. Our data would suggest that during tissue injury, the innate cytokines induced would drive the full activation signal seen, which includes a broad inflammatory response. We note that for example IL17F, which has been recently shown to be pathogenic in IBD models (Tang et al., 2018), is not induced by TCR signals alone but is strongly induced in the presence of inflammatory cytokines (fold change >400, p < 10^−6^).

Overall, we have defined some novel combinatorial pathways to activate MAIT cells, extending the role of TNF superfamily members such as TL1A, and we have dissected the consequences of activation via TCR-dependent and -independent pathways in human blood and gut. Given the overlap between tissue repair and host defence and inflammatory programmes found in MAIT cells (and related populations in the mouse), our model suggests that ongoing maintenance of the barrier is an integral part of the function of such unconventional cells concentrated in epithelia, which goes hand-in-hand with control of microbial invasion.

## MATERIALS AND METHODS

### Samples

All samples were collected with appropriate patient consent and NHS REC provided ethical approval (reference numbers 09/H0606/5 for IBD patients and 16/YH/0247 for controls). Healthy PBMCs were isolated from leukocyte cones (NHS Blood Services). For long-term storage, PBMCs were kept in liquid nitrogen with freezing media (10% DMSO, 90% fetal calf serum, both Sigma-Aldrich). Colonic tissues were collected from uninvolved mucosa of patients with colorectal cancer. Patient information is shown in **Table 1**. All patients involved gave written consent. Colonic tissues were digested at 37°C for overnight with Collagenase A (Roche) and DNase I (Sigma-Aldrich). Colonic lymphocytes were then isolated from the cell suspension using Percoll (GE Healthcare). A detailed protocol has been described by Geremia *et al*. (Geremia et al., 2011).

**Table 1.** Characteristics of the CRC patients.

|  |  |
| --- | --- |
| **CRC patients** | n = 21 |
| **Age** (Average, SD*) | 68, 12 |
| **Sex** (Male/Female) | 12/9 |
| **Uninflamed, tumour-free tissue origin** |  |
| Small intestine | 0 |
| Large intestine | 21 |
| **Time since diagnosis (years)** |  |
| Average, SD | 1, 1 |

\* SD: standard deviation

### Cell culture

PBMCs were thawed, washed and maintained in RPMI 1640 with 10% fetal calf serum, 1% L-glutamine, and 1% penicillin/streptomycin (R10) (all Sigma-Aldrich). CD8^+^ T cells were positively labeled with CD8 Microbeads (Miltenyi Biotech, purities were ≥ 90%), and enriched from PBMCs using MS or LS columns following the manufacturer’s instructions (Miltenyi Biotech). Colonic lymphocytes were used without prior enrichment by CD8 microbeads and were maintained in R10 supplemented with 25ng/mL amphotericin B (Gibco), 40μg/mL gentamicin (Gibco), and 10μg/mL ciprofloxacin (Sigma-Aldrich).

Caco2 cells (Colorectal adenocarcinoma cell line, ATCC) were cultured at a starting density of 4×10^5^ cell/cm^2^ in T-175 cell-culture flasks, using GlutaMAX medium supplemented with 10% fetal bovine serum (FBS), 1% MEM Non-essential-amino acid solution (NEAA), 100ug/mL Penicillin-Streptomycin, 2mM L-glutamine (Sigma Aldrich). Cultures were maintained with media exchange every second day and routinely split every week when cells had reached approximately 70% confluency.

THP1 cells (Human monocyte cell line, ATCC) were cultured at a density between 2×10^5^ to 10^6^ cell/mL in T-175 cell-culture using RPMI-1640 medium (Sigma Aldrich) supplemented with FBS, Penicillin-Streptomycin and L-glutamine.

### *In vitro* stimulations

For non-specific TCR triggering, PBMCs, enriched CD8 T cells, sorted cells or colonic lymphocytes were stimulated with plate bound anti-CD3/28 antibodies (Miltenyi), or anti-CD3/28 beads (Miltenyi) at 1:1 ratio. ELISA plates (Greiner) were coated with 5μg/mL anti-CD3/28 with the final volume of 100μL at 4°C for overnight. Antibody mix was washed off the next morning, and plates were used after 1-hour 37°C incubation with R10. Anti-CD3/28 beads were prepared following the manufacturer’s instructions.

MAIT-specific TCR triggering was achieved by co-culturing of 200,000 MACS-enriched CD8s with 100,000 THP-1 cells which had been previously pulsed with 10nM 5-OP-RU (kindly provided by David Fairlie) for 2 hours. Unpulsed THP-1s were used as controls.

For cytokine triggering, cells were stimulated for 20 hours with IL-12 (Miltenyi) at 2ng/mL, IL-18 (MBL) at 50ng/ml, IL-15 (Miltenyi) at 25ng/ml, TL1A (R&D) at 100ng/ml, unless otherwise stated.

For activation of MAIT cells by bacteria-derived ligands, THP-1 cells were loaded with PFA-fixed (2%, 20min) *E.coli* (DH5α, Invitrogen) at a 25 bacteria per cell (BpC) ratio overnight. Bacterially loaded THP-1s were washed and co-cultured with MACS-enriched CD8s at a 1:2 ratio. In order to block the TCR-dependent component of this activation, in some experiments an anti-MR1 blocking antibody (26.5, Biolegend) was added to the co-cultures.

### Flow cytometry

Brefeldin A (eBioscience, 1000x) was added into the cell culture for the last 4 hours before intracellular staining.

Dyes/human antibodies used were: viability dye Live/Dead fixable-Near-IR (Invitrogen), CD3-eFluor450 (OKT3, eBioscience), CD4-VioGreen (M-T466, Miltenyi), CD4 PerCP-Cyanine5.5 (OKT4, eBioscience), CD8-VioGreen or -Vio770 (REA734, Miltenyi), CD161-PE or -PE-Vio770 (191B8, Miltenyi), Vα7.2-APC or - PE/Cy7 (3C10, BioLegend), Pan Vγδ-FITC (IM1571U, Beckman Coulter), Vγδ-APC-Cy7 (11F2, Miltenyi), IFN-γ-FITC (45-15, Miltenyi), -PE/Cy7 (4S.B3, BioLegend), TNF-α-PerCP-Cy5.5 or -FITC (MAb11, BioLegend), GM-CSF-PerCp/Cy5.5 (BVD2-21C11, Biolegend), CCL3/4-APC (93342, R&D Systems), GrB-APC (GB12, Invitrogen), Furin-AF647 (222722, R&D Systems) CD39-PE (A1, Biolegend), PD-1-BV421 (EH12.2H7, BioLegend), MR1 tetramer-PE (NIH).

Data were acquired on a MACSQuant cytometer (Miltenyi) or LSRII (BD Biosciences) and analyzed on FlowJo (Tree Star Inc.). Gating strategy is shown in **Supplementary Fig. 4A.**

### RNA sequencing (RNAseq)

CD8^+^ T cells were enriched from PBMCs of three healthy individuals and were rested overnight prior to sorting. On the next day, MAIT (CD161^++^ Vα7.2^+^) cells were sorted using a Beckman Coulter MoFlo XDP and stimulated with a range of conditions including anti-CD3/28, cytokines (IL-12/IL-18/IL-15/TL1A), or the combination of both, or left untreated in R10 media for 24 hours. RNA was then extracted from these 12 samples using an RNeasy Micro kit (Qiagen). The quantity and quality of extracted RNA was first evaluated using both a nanodrop spectrophotometer and the Agilent 2100 bioanalyzer. All samples had RNA integrity (RIN) values greater than 9 and were free from contaminating protein and organic compounds. RNAseq was performed by Wellcome Trust Centre for Human Genetics (University of Oxford) on a HiSeq4000v platform. Gene lists that were differentially expressed (>4 fold, P<0.01, FDR<0.05) between various conditions and their normalised expression values, as well as the principle component analysis (PCA) plots, were generated with Partek^®^ Flow^®^, an online analysis platform for Next Generation Sequencing data (http://www.partek.com/partek-flow/), following the user’s guide. Volcano plots were generated with Prism software. Heat maps were generated using normalised counts with Heat Mapper (http://www.heatmapper.ca/expression/) with the averaged linkage clustering method and Pearson distance measurement method. Venn diagrams were drawn with an online diagram drawing platform developed by Ghent University, Belgium (http://bioinformatics.psb.ugent.be/webtools/Venn/). Gene set enrichment analysis (GSEA) was performed using GSEA version 3.0 (Subramanian et al., 2005), comparing gene expression data as a whole with the reference gene list obtained from the publication by Linehan *et al*. (Linehan et al., 2018).

### qPCR

CD161^++^Vα7.2^+^ (MAIT) and CD161^−^Vα7.2^−^ cells were sorted from pre-enriched blood CD8^+^ T cells. These cells were then stimulated with anti-CD3/28, cytokines (IL-12/IL-18/IL-15/TL1A), or the combination of both, or left untreated in R10 media for 20 hours. The total RNA of sorted T cells was extracted with an RNeasy Micro kit (Qiagen) and reverse transcribed using reverse transcribed using SuperScript III Reverse Transcriptase (Invitrogen). For detection of IL-26 mRNA, a 20X IL-26 human TaqMan^®^ probe was used (Hs00218189_m1) with 2X TaqMan^®^ Fast Advanced Master Mix (both from ThermoFisher Scientific). OSM and HEBGF cDNA quantification was performed with Roche ^®^ hydrolysis probes (*OSM*: forward primer sequence, 5’-cttccccagtgaggagacc-3’, reverse sequence, 5’-ctgctctaagtcggccagtc-3’; HBEGF: forward primer sequence, 5’-tggggcttctcatgtttagg-3’, reverse sequence, 5’-catgcccaacttcactttctc-3’), with GAPDH as the internal control (forward primer sequence, 5’-ccccggtttctataaattgagc-3’, reverse sequence, 5’-cttccccatggtgtctgag-3’).

### In vitro wound-healing assay

Enriched CD8^+^ were co-cultured with THP1 cells loaded with fixed *E.coli* at 30 BpC in the presence or absence of 20ug/mL LEAF™ anti-MR1 Antibody (Biolegend). Supernatants were collected at 72 hours. A total of 1.5×10^4^ Caco2 cells were seeded per well in a 96-well clear flat bottom plate (Corning) and grown to confluency at 37C for 5 days with media exchange every 2 days. Monolayers were scratched using a WoundMaker (Essen Bioscience), washed with serum-free medium and incubated with CD8^+^ 72h hour supernatants diluted 1:4 with fresh media. As a negative control, fresh media was used. Time lapse imaging was recorded every 4 hours using IncuCyte S3 Live Cell Analysis System (Essen Bioscience) for 36 hours at 37°C. GlutaMAX medium supplemented with 10% FBS, 1% NEAA, Penicillin-Streptomycin and L-glutamine was used throughout this experiment.

### Data analysis

All graphs and statistical analyses, except RNAseq data analysis, were completed using GraphPad Prism Software Version 6.0b (La Jolla, CA). Statistical significance was assessed using paired Student’s t-test, or repeated-measures two-way analysis of variances, with Bonferroni’s correction for multiple comparison assays. For the invitro wound-healing assay, ImageJ v1.8 was used to determine the area of wounds. Area at different timepoints were normalised as a percentage of the initial area. All data were presented as means with s.e.m.

## SUPPLEMENTAL INFORMATION

Supplemental Information includes four figures and five tables (tables can be found in :https://www.dropbox.com/sh/ck27pfu8e4a6yn0/AABgjixZIkop2mGDuvea5yW2a?dl=0).

SFigure 1 (related to Fig 1). **TL1A and IL-15 alone do not promote MAIT cell effector functions and have only a limited effect on CD161+ and CD161-CD8+ T-cells.** CD8+ T cells were enriched from healthy PBMCs and stimulated overnight with combinations of the indicated cytokines. **(A)** Proportion of CD8+ MAIT cells producing CD69, IFN-γ, and TNF-αwhen left untreated, or stimulated singly with 100ng/ml TL1A. **(B-D)** Frequency of MAIT cells expressing IFN-γ (B), TNF-α (C) or Grb (D) upon stimulation with TL1A (100ng/ml), IL-15 (25ng/ml) or both cytokines. **(E)** Gating strategy for CD8+ MAIT (CD161++Vα7.2+)/CD161+Vα7.2-/CD161-Vα7.2-cells **(F)** Proportion of CD8+MAIT (CD161++Vα7.2+), CD161+Vα7.2-, or CD161-Vα7.2-cells producing IFN-γ when stimulated with IL-12, IL-15, IL-18 and TL1A. **(G)** Proportion of CD8+CD161+Vα7.2-cells producing IFN-γ when stimulated with a range of conditions. **(H-M)** Proportion of CD8+CD161+Vα7.2-cells (H-J) or CD8+CD161-Vα7.2-cells (K-M) producing IFN-γ, TNF-α, or GrB when treated with combinations of TL1A (100ng/ml) and IL-15 (25ng/ml) with suboptimal IL-12/18 (2ng/ml). Data were acquired from 6-8 donors in 2-3 experiments. Differences between the conditions were analysed by Friedman tests with Dunn’s multiple comparison tests.

SFigure 2 (related to Fig. 2). **Functional studies on the impact of combined TCR and cytokine signalling**. CD8+ T cells were enriched from healthy PBMCs and stimulated in different ways. **(A, B)** Proportion of CD8+ MAIT (CD161 ++Vα7.2+)/CD161+/CD161 - cells producing IFN-γ (A) or TNF-α (B) following overnight incubation with suboptimal concentrations of IL-12 and IL-18, plus 〈CD3/CD28 beads at increasing bead-to-cell ratios. **(C)** Proportion of CD8+MAIT cells producing IFN-γ (following stimulation with increasing concentrations of cytokines: IL-12, IL-18, or TL1A, respectively in the presence of plate-bound αCD3/CD28 antibodies. Data were acquired from 7-8 donors in three experiments. Differences between the conditions were analysed by 2way ANOVA with Tuckeys multiple comparison tests (A-C). ns = not significant, **p<0.01, ***p<0.001, ****p<0.0001.

SFigure 3 (related to Fig. 3). **Identification of MAIT cells in the colonic lamina propria and additional functional studies on blood-derived MAITs. (A)** Gating strategy to identify CD8+MAIT cells from gut LPLs. **(B)** Proportion of CD8+MAIT cells producing IFN-γ or TNF-α after overnight stimulations. CD8+MAIT cells were derived from PBMCs, which, prior to stimulation, were either rested in the normal media or stirred in the digestion media containing DNase and Collagenase A for 12 hours. **(C)** Representative plot showing how to identify MAIT cells from the gut by using either a conventional Vα7.2 TCR staining antibody or the MR1-tetramer staining antibody, in combination with CD161 staining. **(D)** Proportion of CD8+ MAIT cell expressing the indicated molecules after overnight co-culture with THP1 cells incubated with 25 fixed *E.coli* bacteria per cell in the presence of an blocking antibody directed against MR1 or an isotype control. Data were acquired from 1-7 donors in 1-3 experiments. Differences between the conditions were analysed by Wilcoxon tests (D). *p<0.05

SFigure 4 (related to Fig. 5). **Expression of effector molecules by MAITs treated with the conditions used in the RNAseq study.** CD8+ T-cells were MACS enriched and left untreated (UT) or were stimulated with αCD3/28 beads (T), suboptimal IL-12/18 in combination with TL1A and IL-15 (C) or with a combination of the aforementioned cytokines and αCD3/28 beads (TC) overnight. **(A-D)** Proportion of CD8+MAIT cells isolated from parts of the samples used for the RNAseq experiment producing IFN-γ (A), TNF-α (B), GrB (C) or CD69 (D). Each dot corresponds to a donor of the RNAseq study, data were acquired from 3 donors in one experiment. **(E-F)** Expression levels of HBEGF (E), OSM (F) and IL26 (G) in CD8+MAIT cells (n=5) examined by qPCR. GAPDH was used as house-keeping gene. Data were acquired from five donors in two experiments. Differences between conditions were analysed by Friedman tests with Dunn’s multiple comparisons tests. *p<0.05, **p<0.01.

SFigure 5 (related to Fig. 6). **Further investigation of tissue repair related functions of MAIT cells. (A, B)** Relative expression of the genes of the tissue repair gene set by MAIT cells stimulated by TCR (A) or TCR+cytokines (B) compared to unstimulated controls. The leading edge genes of the corresponding GSEA plots (F6) are marked. The original tissue repair gene set of 101 genes was restricted to the genes present in our dataset. Data were acquired from 3 donors in one experiment. **(C)** Flow cytometry analysis of the expression of TNF-α, Furin and CCL3 by CD161++/MAIT CD8+ T cells in response to fixed *E.coli* presented by THP1 cells in the presence or absence of an anti-MR1 (aMR1) blocking antibody at 20h timepoint. **(D)** Statistical analysis of the expression of the effector molecules shown in (A). Data were acquired from seven donors in three experiments. **(E, F)** Flow cytometry and statistical analysis of the expression of GM-CSF by CD161++/MAIT CD8+ T cells at 20h and 72h timepoints using the conditions described in (A). Data were acquired from three donors in one experiment.

## ACKNOWLEDGEMENTS

This research was supported by the Wellcome Trust (WT109965MA), NIHR Senior Fellowship (PK), NIHR Biomedical Research Centre, Oxford, NIH U19 I082360, Chinese Scholarship Council (TL). The authors wish to acknowledge the BRC Oxford GI Biobank and BRC Oxford IBD Cohort in collecting and making available the samples/data used in the generation of this publication. In particular, we thank Ahmed Hegazy, Nathaniel West, James Chivenga, Cloe Vassart, and David Maldonado-Perez for their effort and help. We additionally thank Chan Phetsouphanh for sorting the cells in the Peter Medawar Building.

5-OP-RU was a kind gift from David Fairlie, University of Queensland.

## AUTHOR CONTRIBUTIONS

Conceived and designed the experiments: T.L., H.A., C-P.H., V.M., P.K., C.W., D.E.;

Performed the experiments: T.L., H.A., C-P.H, V.M., T.K.;

Analyzed the data: T.L., H.A., C-P.H., T.K., P.K.;

Contributed to the collection of clinical samples: T.L., T.K., M.F., Z.C., S.M., N.W., M.N., R.P.;

Contributed to the writing of this manuscript: T.L., H.A., C-P.H., P.K., C.W.

