## Supplemental figures and tables for "TCR and inflammatory signals tune human MAIT cells to exert specific tissue repair and effector functions"

**KEY RESOURCES TABLE**

| REAGENT or RESOURCE | SOURCE | IDENTIFIER |
|---|---|---|
| Antibodies | | |
| LEAF-purified mouse anti-human CD3 (clone UCHT1) | Biolegend | Discontinued Alternative: Cat# 300438 |
| LEAF-purified mouse anti-human CD28 (clone CD28.2) | Biolegend | Cat# 302914 |
| LEAF-purified mouse anti mouse/rat/human MR1 (clone 26.5) | Biolegend | Cat# 361103 |
| Anti-human CD3 (clone OKT3) eFluor450 | eBioScience | Cat# 48-0037-42 |
| Anti-human CD3 (clone OKT3) BV605 | Biolegend | Cat# 317321 |
| Anti-human CD4 (clone M-T466) VioGreen | Miltenyi | Cat# 130-113-259 |
| Anti-human CD4 (clone OKT4) PerCP/Cy5.5 | eBioScience | Cat# 45-0048-42 |
| Anti-human CD8 (clone REA734) VioGreen | Miltenyi | Cat# 130-110-684 |
| Anti-human CD8 (clone REA734) PE-Vio770 | Miltenyi | Cat# 130-110-680 |
| Anti-human CD39 (clone A1) PE | Biolegend | Cat# 328207 |
| Anti-human CD69 (clone H1.2F3) eFluor450 | eBioScience | Cat# 48-0691-82 |
| Anti-human CD161 (clone 191B8) APC | Miltenyi | Cat# 130-113-590 |
| Anti-human CD161 (clone 191B8) PE | Miltenyi | Cat# 130-113-593 |
| Anti-human CD161 (clone 191B8) PE-Vio770 | Miltenyi | Cat# 130-113-594 |
| Anti-human CCL3/(4) (clone 93342) APC | R&D Systems | Cat# AF270NA |
| Anti-human Furin (clone 222722) AF647 | R&D Systems | Cat# IC1503R-100UG |
| Anti-human GM-CSF (clone BVD2-21C11) PerCP/Cy5.5 | Biolegend | Cat# 502312 |
| Anti-human GrB (clone GB12) APC | Initrogen | Cat# MHGB05 |
| Anti-human GrB (clone GB11) AF700 | BD BioSciences | Cat# 561016 |
| Anti-human IFNγ (clone 4S.B3) AF700 | Biolegend | Cat# 502520 |
| Anti-human IFNγ (clone 45-15) FITC | Miltenyi | Cat# 130-091-641 |
| Anti-human IFNγ (clone 4S.B3) PE/Cy7 | Biolegend | Cat# 502528 |
| Anti-human IgG2bκ (clone 133303) AF647 | R&D Systems | Cat# IC0041R |
| Anti-human PD-1 (clone EH12.2H7) BV421 | Biolegend | Cat# 329920 |
| Anti-human TNFα (clone Mab11) FITC | Biolegend | Cat# 502906 |
| Anti-human TNFα (clone Mab11) PerCP/Cy5.5 | Biolegend | Cat# 502926 |
| Anti-human Vα7.2 (clone 3C10) APC | Biolegend | Cat# 351708 |
| Anti-human Vα7.2 (clone 3C10) FITC | Biolegend | Cat# 351704 |
| Anti-human Vα7.2 (clone 3C10) PE/Cy7 | Biolegend | Cat# 31712 |
| Anti-human TCRγ/δ (clone IMMU510) FITC | Beckman Coulter | Cat# IM1571U |
| Anti-human TCRγ/δ (clone 11F2) APC-Vio770 | Miltenyi | Cat# 130-113-501 |
| Bacterial Strains | | |
| *E.coli* DH5α | Invitrogen | Cat# 18265017 |
| Biological Samples | | |
| Leukocyte cones | NHS Blood and Transplant | https://www.nhsbt.nhs.uk/ |
| Patient-derived resections (CRC) | TGU Biobank | https://www.expmedndm.ox.ac.uk/tgu/tgu-biobank-ibd-cohort |
| Patient-derived resection (IBD) | TGU Biobank | https://www.expmedndm.ox.ac.uk/tgu/tgu-biobank-ibd-cohort |

| | | |
|---|---|---|
| **Chemicals, Peptides, and Recombinant Proteins** | | |
| Brefeldin A solution (1000X) | eBioScience | Cat# 00-4506-51 |
| 5-OP-RU | Fairlie group<br>Mak et al. 2017 | N/A |
| Collagenase A | Roche (mft)/ Merck | Cat# 10103578001 |
| DNase I | Roche (mft)/ Merck | Cat# 11284932001 |
| SuperScript III Reverse Transcriptase | Invitrogen | Cat# 18080093 |
| Human IL-12, premium grade | Miltenyi | Cat# 130-096-705 |
| Human IL-15, premium grade | Miltenyi | Cat# 130-095-764 |
| Recombinant Human IL-18 | MBL | Cat# B001-5 |
| Recombinant Human TL1A/TNFSF15 | R&D Systems | Cat# 1319-TL-010 |
| **Critical Commercial Assays** | | |
| T cell Activation/Expansion Kit, human | Miltenyi | Cat# 130-091-441 |
| RNeasy Micro Kit | Quiagen | Cat# 74004 |
| LIVE/DEAD Fixable Near IR Dead Cell Stain Kit | Invitrogen | Cat# L10119 |
| Permeabilization buffer (10x) | eBioScience | Cat# 00-8333-56 |
| CD8 MicroBeads, human | Miltenyi | Cat# 130-045-201 |
| **Deposited Data** | | |
| RNA-seq files (UT, T, C, CT) | GEO | GSE129906 |
| **Experimental Models: Cell Lines** | | |
| THP-1 | ATCC | TIB-202 |
| Caco2 | ATCC | HTB-37 |
| **Oligonucleotides** | | |
| Primer specific for *OSM*: Forward > cttccccagtgaggagacc | Roche | N/A |
| Primer specific for *OSM*: Reverse > ctgctctaagtcggccagtc | Roche | N/A |
| Primer specific for *HBEGF*: Forward > tggggcttctcatgtttagg | Roche | N/A |
| Primer specific for *HBEGF*: Reverse > catgcccaacttcactttctc | Roche | N/A |
| Primer specific for *GAPDH*: Forward > ccccggtttctataaattgagc | Roche | N/A |
| Primer specific for *GAPDH*: Reverse> cttccccatggtgtctgag | Roche | N/A |
| **Software and Algorithms** | | |
| Bioinformatics & Evolutionary Genomics | Ghent University | http://bioinformatics.psb.ugent.be/webtools/Venn/ |
| FlowJo 10 | Tree Star | https://www.flowjo.com |
| GSEA version 3.0 | Subramanian et al. 2005 | https://software.broadinstitute.org/gsea/index.jsp |
| Heatmapper | Wishart group,<br>University of Alberta | http://www.heatmapper.ca |
| Image J Version 1.8 | NIH | https://imagej.nih.gov/ij/index.html |
| Partek Flow | Partek | http://www.partek.com/partek-flow/ |
| Prism Version 6.0b | Graphpad | https://www.graphpad.com |

| Other | | |
|---|---|---|
| 5-OP-RU-MR1-Tetramer PE | NIH tetramer core facility | N/A |
| PerColl | GE Healthcare (mft)/ Merck | Cat# GE17-0891-01 |
| Human *IL26* TaqMan Probe | Thermo Fisher Scientific | Hs00218189_m1 |
| TaqMan™ Fast Advanced Master Mix | Thermo Fisher Scientific | Cat# 4444557 |

Supplemental Figures

### Supplementary Figure 1

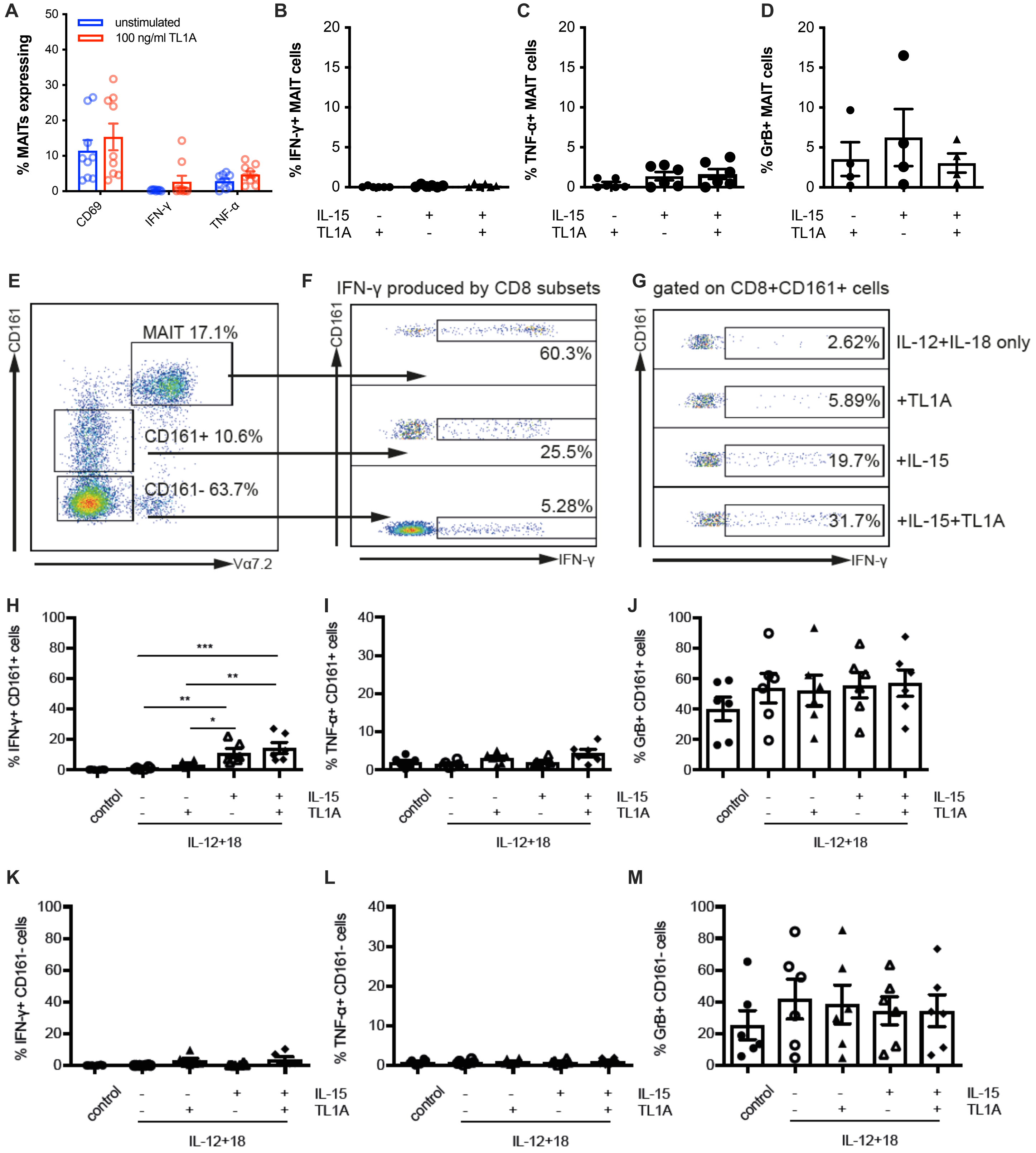


SFigure 1. **TL1A and IL-15 alone do not promote MAIT cell effector functions and have only a limited effect on CD161+ and CD161- CD8+ T-cells.** CD8+ T cells were enriched from healthy PBMCs and stimulated overnight with combinations of the indicated cytokines. **(A)** Proportion of CD8+ MAIT cells producing CD69, IFN-γ, and TNF-α when left untreated, or stimulated singly with 100ng/ml TL1A. **(B-D)** Frequency of MAIT cells expressing IFN-γ (B), TNF-α (C) or Grb (D) upon stimulation with TL1A (100ng/ml), IL-15 (25ng/ml) or both cytokines. **(E)** Gating strategy for CD8+ MAIT (CD161++Vα7.2+)/CD161+Vα7.2-/CD161-Vα7.2-cells **(F)** Proportion of CD8+MAIT (CD161++Vα7.2+), CD161+Vα7.2-, or CD161-Vα7.2-cells producing IFN-γ when stimulated with IL-12, IL-15, IL-18 and TL1A. **(G)** Proportion of CD8+CD161+Vα7.2-cells producing IFN-γ when stimulated with a range of conditions. **(H-M)** Proportion of CD8+CD161+Vα7.2- cells (H-J) or CD8+CD161-Vα7.2- cells (K-M) producing IFN-γ, TNF-α, or GrB when treated with combinations of TL1A (100ng/ml) and IL-15 (25ng/ml) with supoptimal IL-12/18 (2ng/ml). Differences between the conditions were analysed by Friedman tests with Dunn's multiple comparision tests.

### Supplementary Figure 2

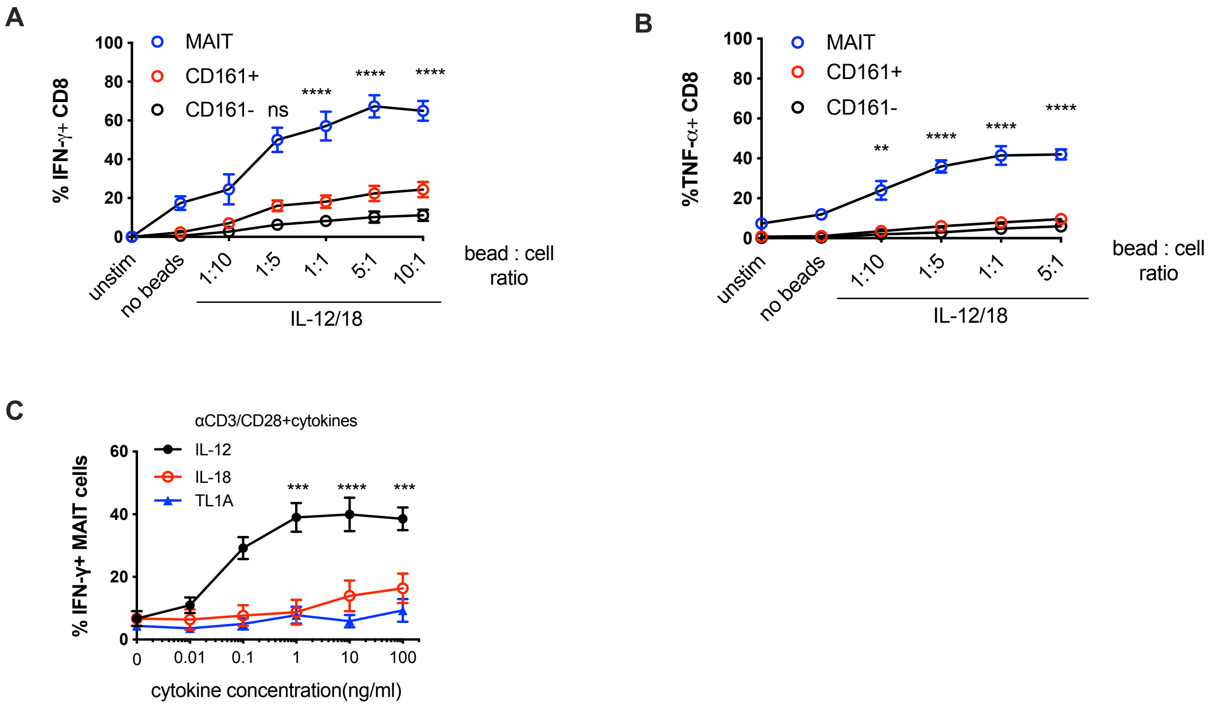


SFigure 2. **Functional studies on the impact of combined TCR and cytokine signalling.** CD8+ T cells were enriched from healthy PBMCs and stimulated in different ways. **(A, B)** Proportion of CD8+ MAIT (CD161++Vα7.2+)/ CD161+/CD161- cells (n=8) producing IFN-γ (A) or TNF-α (B) following overnight incubation with suboptimal concentrations of IL-12 and IL-18, plus αCD3/CD28 beads at increasing bead-to-cell ratios. **(C)** Proportion of CD8+MAIT cells producing IFN-γ following stimulation with increasing concentrations of cytokines: IL-12 (n=8), IL-18 (n=8), or TL1A (n=7), respectively in the presence of plate-bound αCD3/CD28 antibodies. Differences between the conditions were analysed by 2way ANOVA with Tuckeys multiple comparision tests (A-C). ns = not significant, **p<0.01, ***p<0.001, ****p<0.0001.

**Supplementary Figure 3**

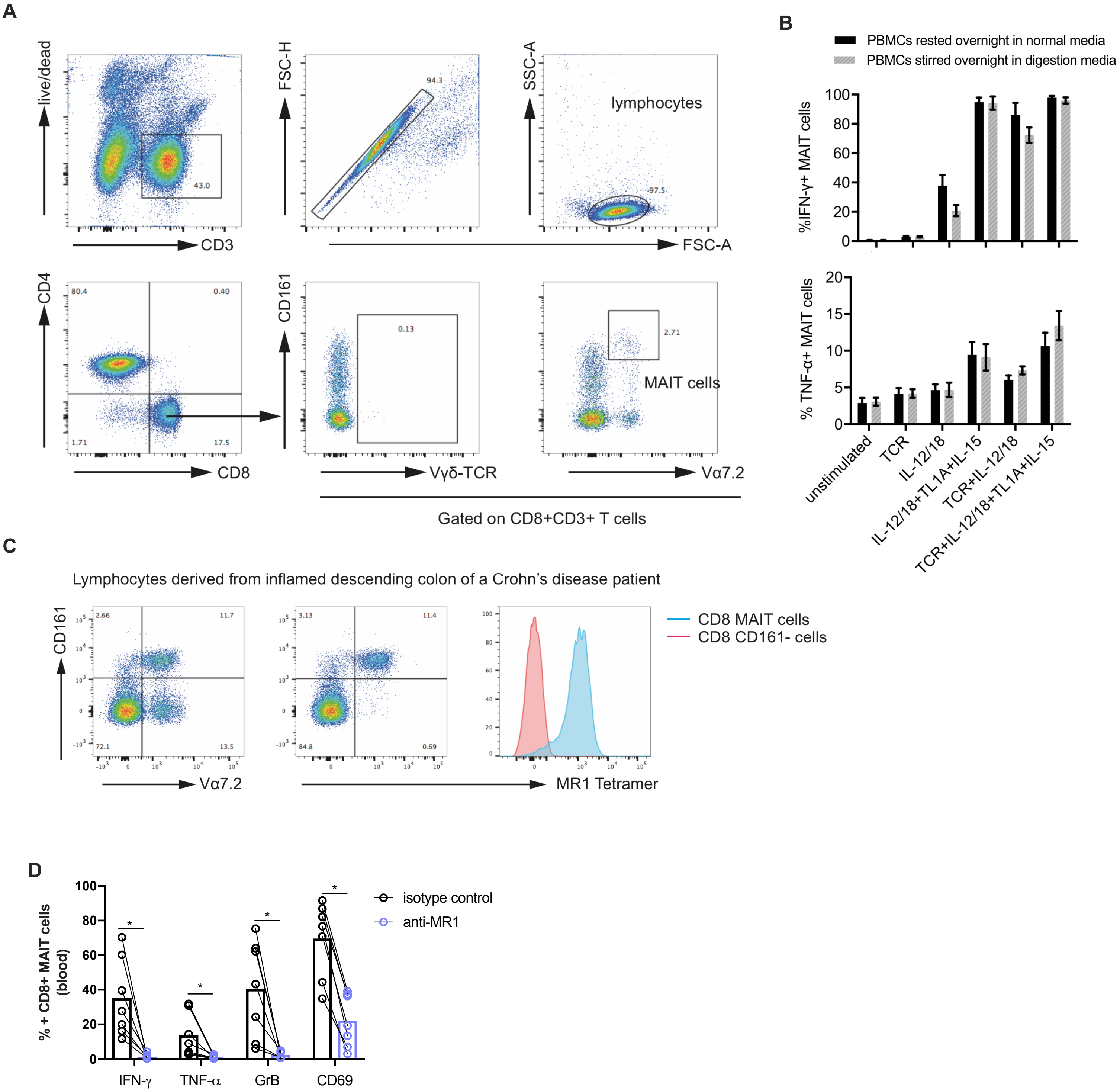


SFigure 3. **Identification of MAIT cells in the colonic lamina propria and additonal functional studies on blood-derived MAITs. (A)** Gating strategy to identify CD8+MAIT cells from gut LPLs. **(B)** Proportion of CD8+MAIT cells producing IFN-γ or TNF-α after overnight stimulations. CD8+MAIT cells were derived from PBMCs, which, prior to stimulation, were either rested in the normal media or stirred in the digestion media containing DNase and Collagenase A for 12 hours. (**C)** Representative plot showing how to identify MAIT cells from the gut by using either a conventional Vα7.2 TCR staining antibody or the MR1-tetramer staining antibody, in combination with CD161 staining. **(D)** Proportion of CD8+ MAIT cell expressiong the indicated molecules after overnight co-culture with THP1 cells incubated with 25 fixed *E.coli* bacteria per cell in the presence of an blocking antibidy directed against MR1 or an isotype control. Differences between the conditions were analysed br Wilcoxon tests (D). *p<0.05

**Supplementary Figure 4**

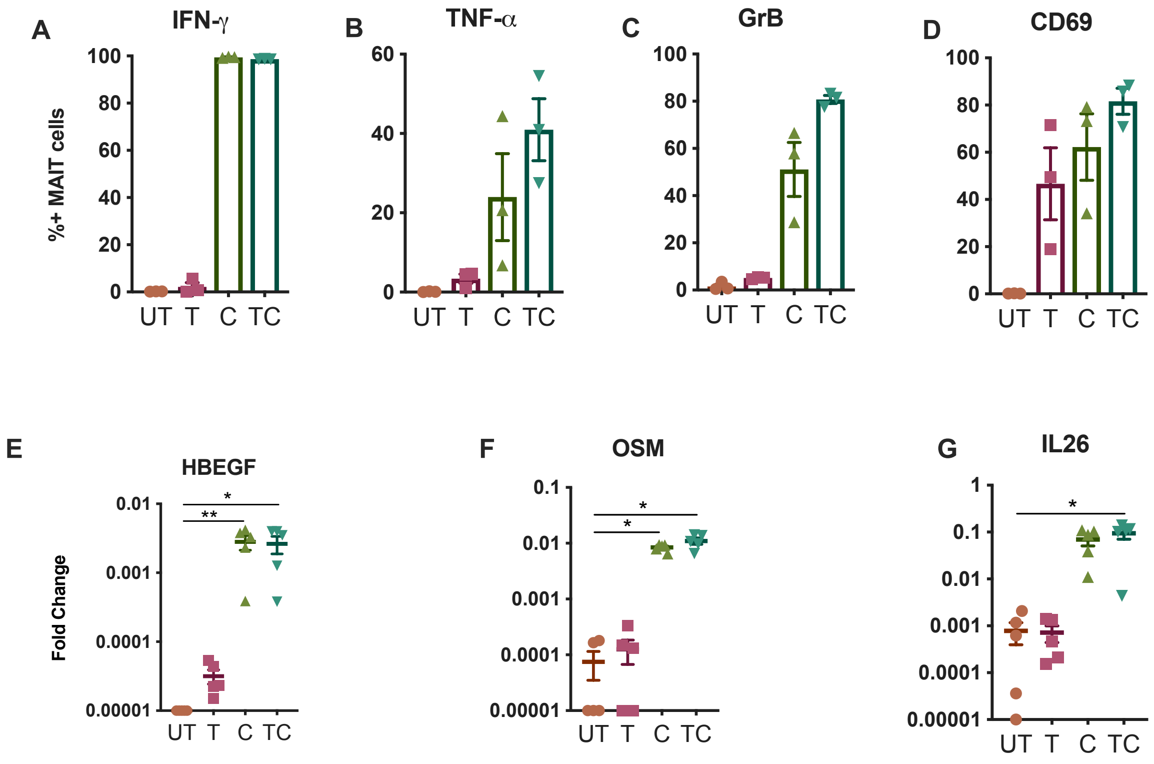


SFigure 4. **Expression of effector molecules by MAITs treated with the conditions used in the RNAseq study.** CD8+ T-cells were MACS enriched and left untreated (UT) or were stimulated with αCD3/28 beads (T), suboptimal IL-12/18 in combination with TL1A and IL-15 (C) or with a combination of the aforementioned cytokines and αCD3/28 beads (TC) overnight. **(A-D)** Proportion of CD8+MAIT cells isolated from parts of the samples used for the RNAseq experiment producing IFN-γ (A), TNF-α (B), GrB (C) or CD69 (D). Each dot corresponds to a donor of the RNAseq study. **(E-F)** Expression levels of HBEGF (E), OSM (F) and IL26 (G) in CD8+MAIT cells (n=5) examined by qPCR. GAPDH was used as house-keeping gene. Data shown are pooled from twoindividual experiments.Differences between conditions were analysed by Friedman tests with Dunn's multiple comparisons tests. *p<0.05,**p<0.01.

Suplementary Figure 5

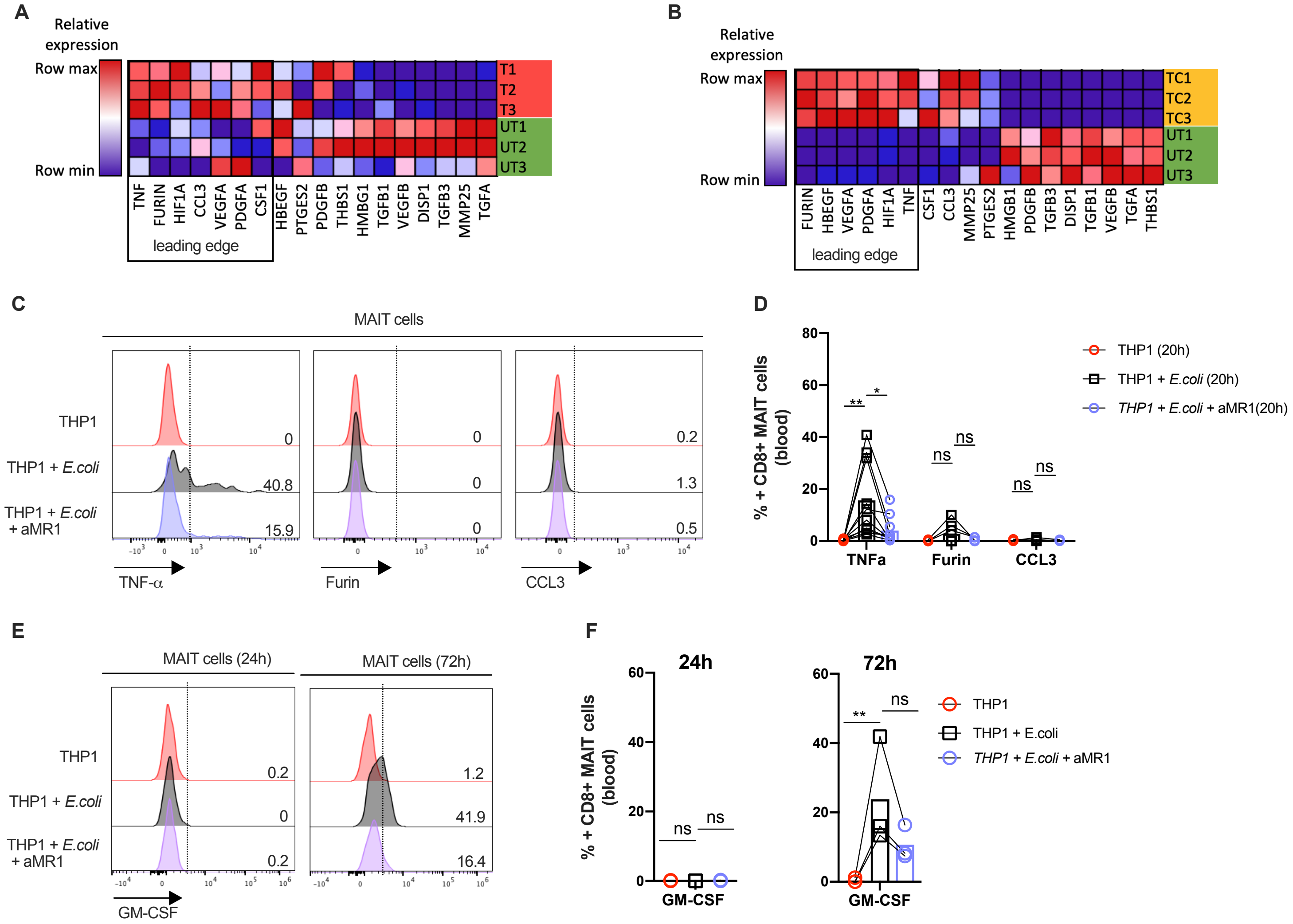

Click here to access/download
**Supplemental Videos and Spreadsheets**
Supplementary table 1.xlsx

Click here to access/download
**Supplemental Videos and Spreadsheets**
Supplementary table 2.xlsx

Supplemental Spreadsheets

Click here to access/download
**Supplemental Videos and Spreadsheets**
Supplementary table 3.xlsx
